## Supplemental figures for "Bark beetles locate fungal symbionts by detecting volatile fungal metabolites of host tree resin monoterpenes"

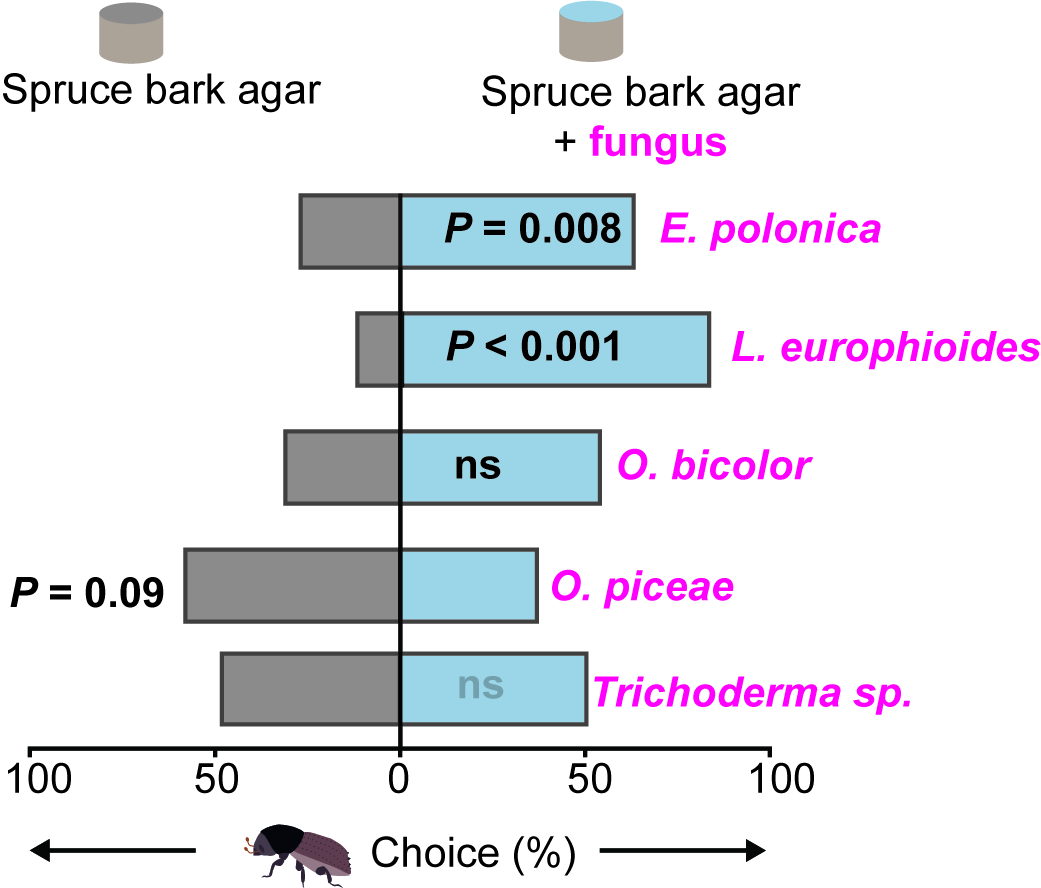

**Figure S1**: Adult beetles prefer spruce bark agar inoculated with two species of symbiotic fungi over uninoculated SBA. Adult beetles did not prefer *O. bicolor*, *O. piceae*, and *Trichoderma sp.*,the latter two species are saprophytes. Deviation of response indices against zero was tested using Wilcoxon’s test. Asterisks denotes significant differences, **P* < 0.05, ***P* < 0.01, ****P* < 0.001 (*n = 25*).

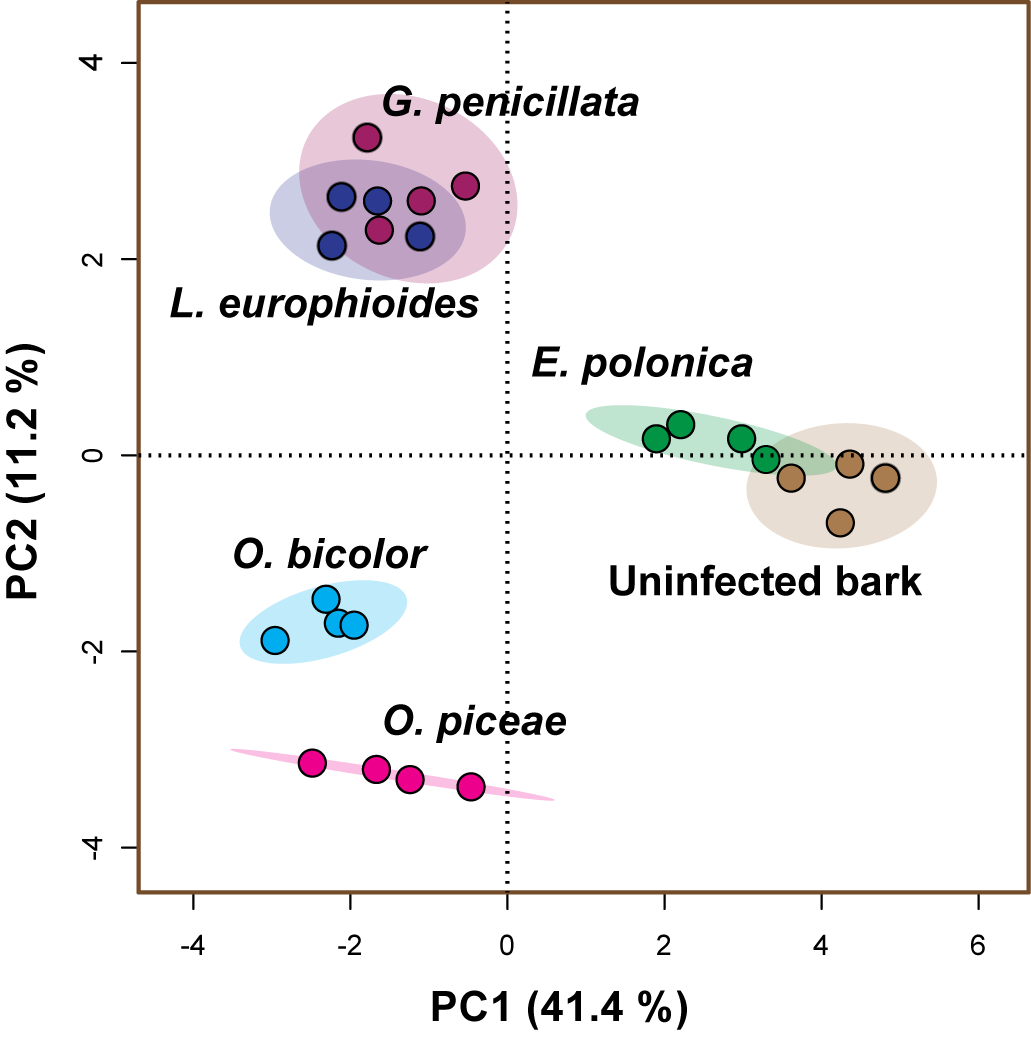

**Figure S2:** Volatile emission pattern differed between spruce bark inoculated with different fungi and uninfected bark 4 days after inoculation, as depicted in a sparse [partial least squares discriminant analysis (sPLS-DA)](https://www.metaboanalyst.ca/MetaboAnalyst/Secure/analysis/AnalysisView.xhtml). Principal components (PC1 and PC2) explain 41.4% and 11.2% of the total variation, respectively, and ellipses denote 95% confident intervals around each species. The sPLS-DA plot was generated by using MetaboAnalyst 3.0 software with normalized data (both log transformed and range scaled).

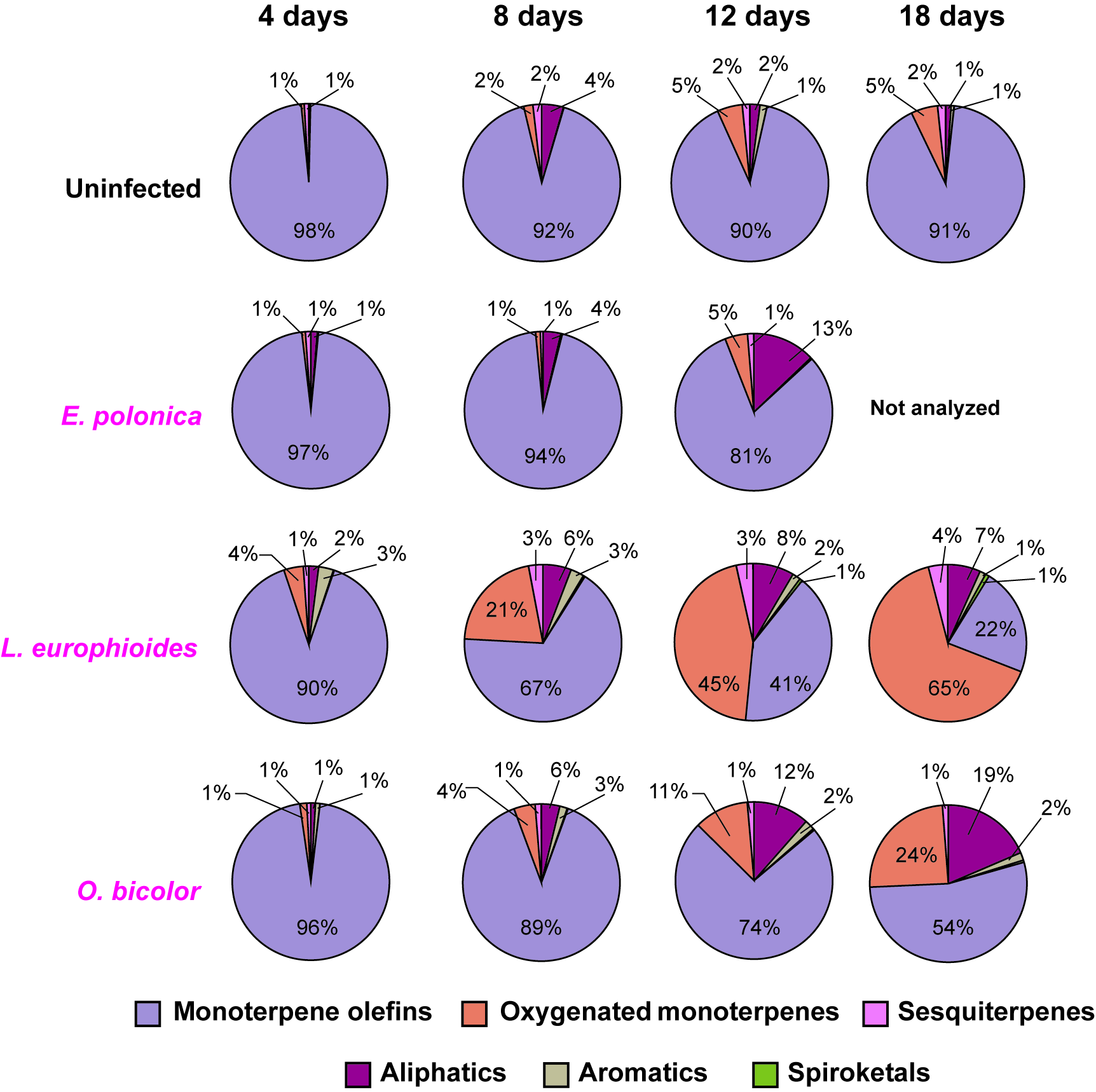

**Figure S3**: Changes in volatile emission profiles of fungal-infested vs. uninfested spruce bark over an 18 day time course for three other *I. typographus* symbiotic fungi besides *G. penicillata*. Compounds are classified into six groups according to chemical structures. Complete volatile emission data by compound and time point for each fungal species are given in Tables S2-S6. (*n* = 5).

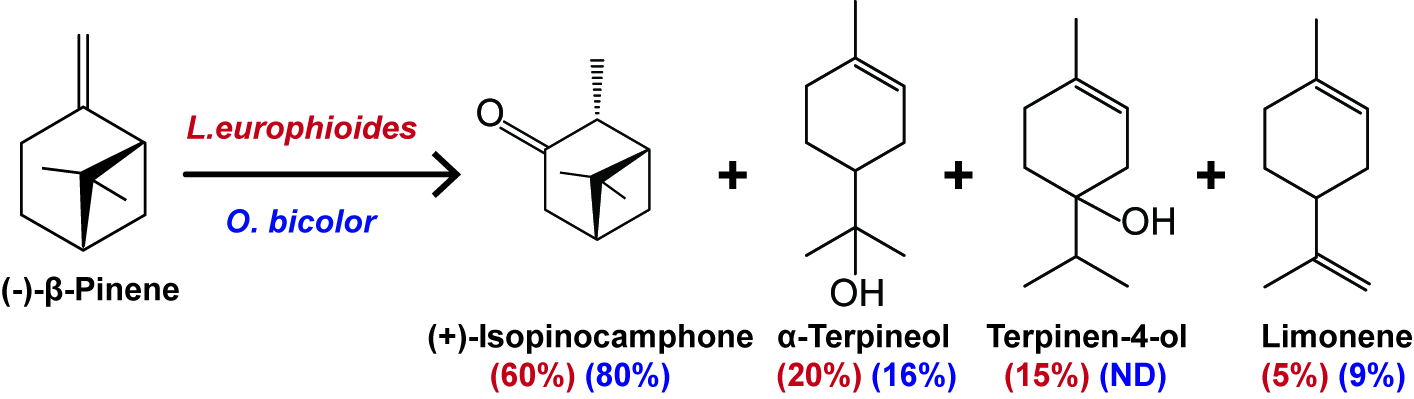

**Figure S4**: Volatile metabolites of (-)-β-pinene produced by two fungal symbionts (*L. europhioides* and *O. bicolor*) of *I. typographus* growing on potato dextrose agar. Isopinocamphone was the major biotransformation product (*n* = 5). *Endoconidiophora polonica* produced no detectable products. ND=not detected.

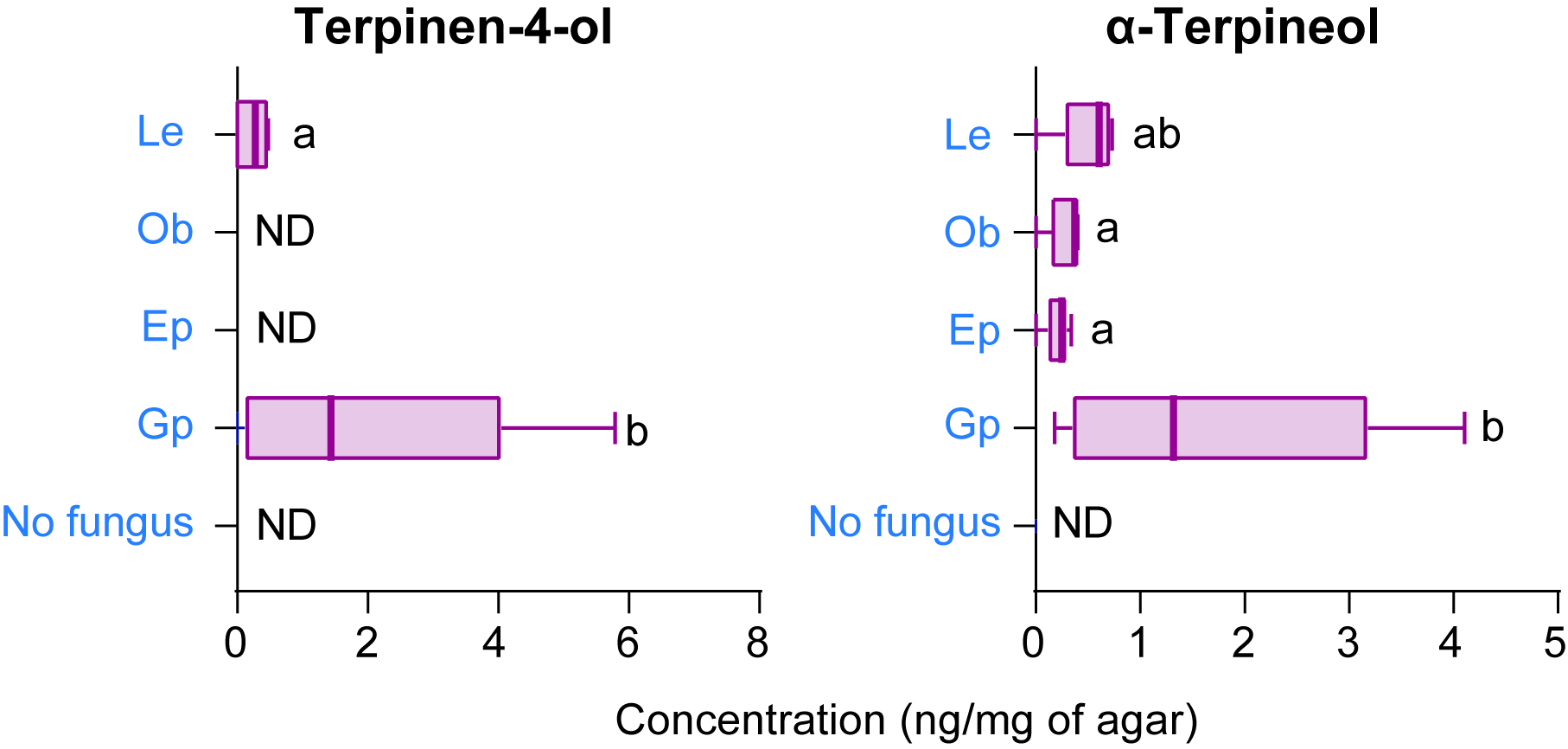

**Figure S5**: Metabolites of (-)-β-pinene produced by various *I. typographus* symbiotic fungi growing on potato dextrose agar after this monoterpene was administered to cultures of each species. Amounts of metabolites were determined after hexane extraction of the agar. Error bars represent SEM (*n* = 5). ND=not detected. Different lowercase letters denote significant differences between treatments (ANOVA, Sidak’s test; *P<*0.05). Fungal abbreviations: *E. polonica* (Ep), *L. europhioides* (Le), *G. penicillata* (Gp), *O. bicolor* (Ob).

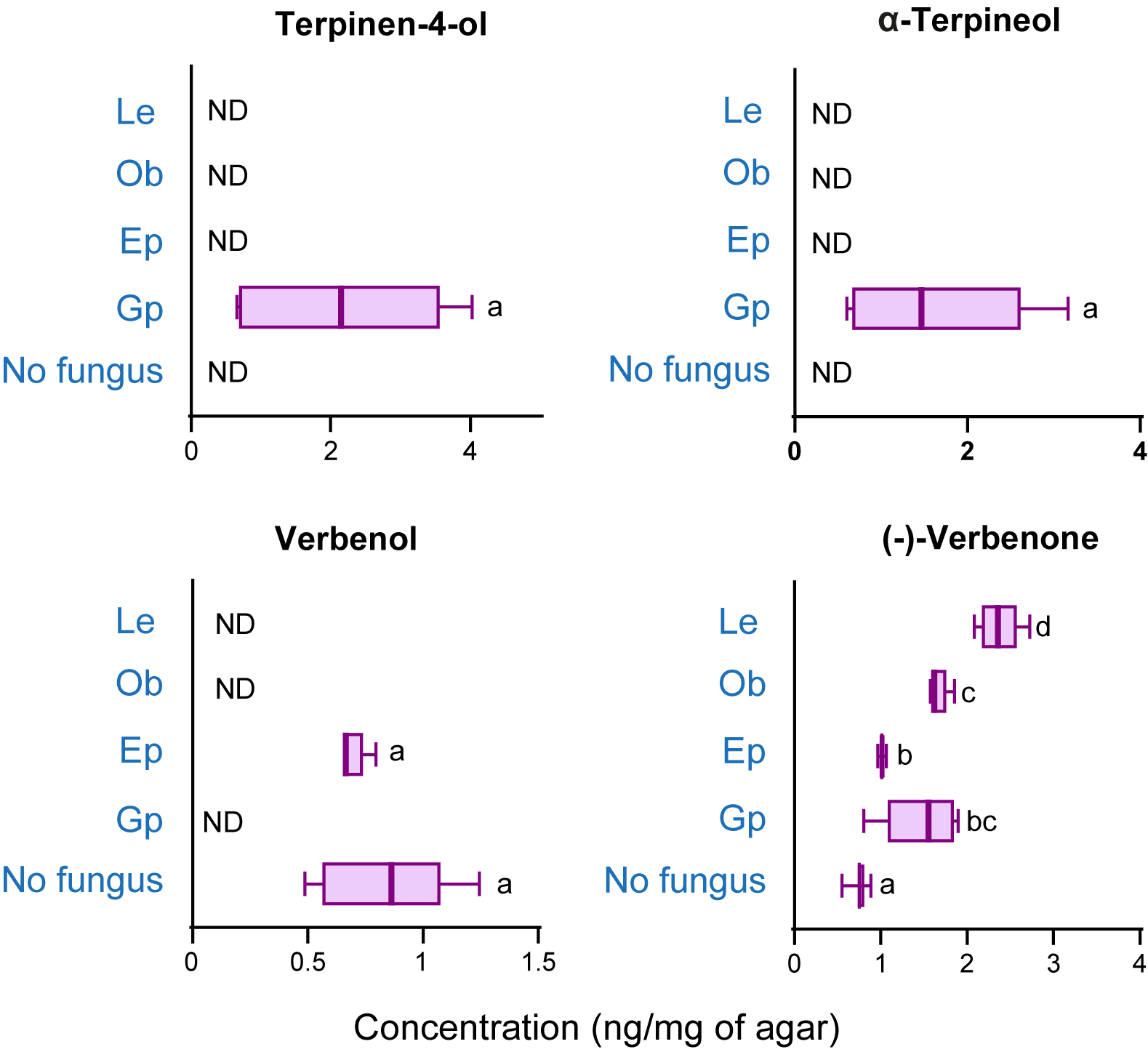

**Figure S6**: Metabolites of (-)-α-pinene produced by various *I. typographus* symbiotic fungi growing on potato dextrose agar after this monoterpene was administered to cultures of each species. Amounts of metabolites were determined after hexane extraction of the agar. Error bars represent SEM (*n* = 5). ND=not detected. Different lowercase letters denote significant differences between treatments (ANOVA, Sidak’s test; *P<*0.05). Fungal abbreviations: *E. polonica* (Ep), *L. europhioides* (Le), *G. penicillata* (Gp), *O. bicolor* (Ob).

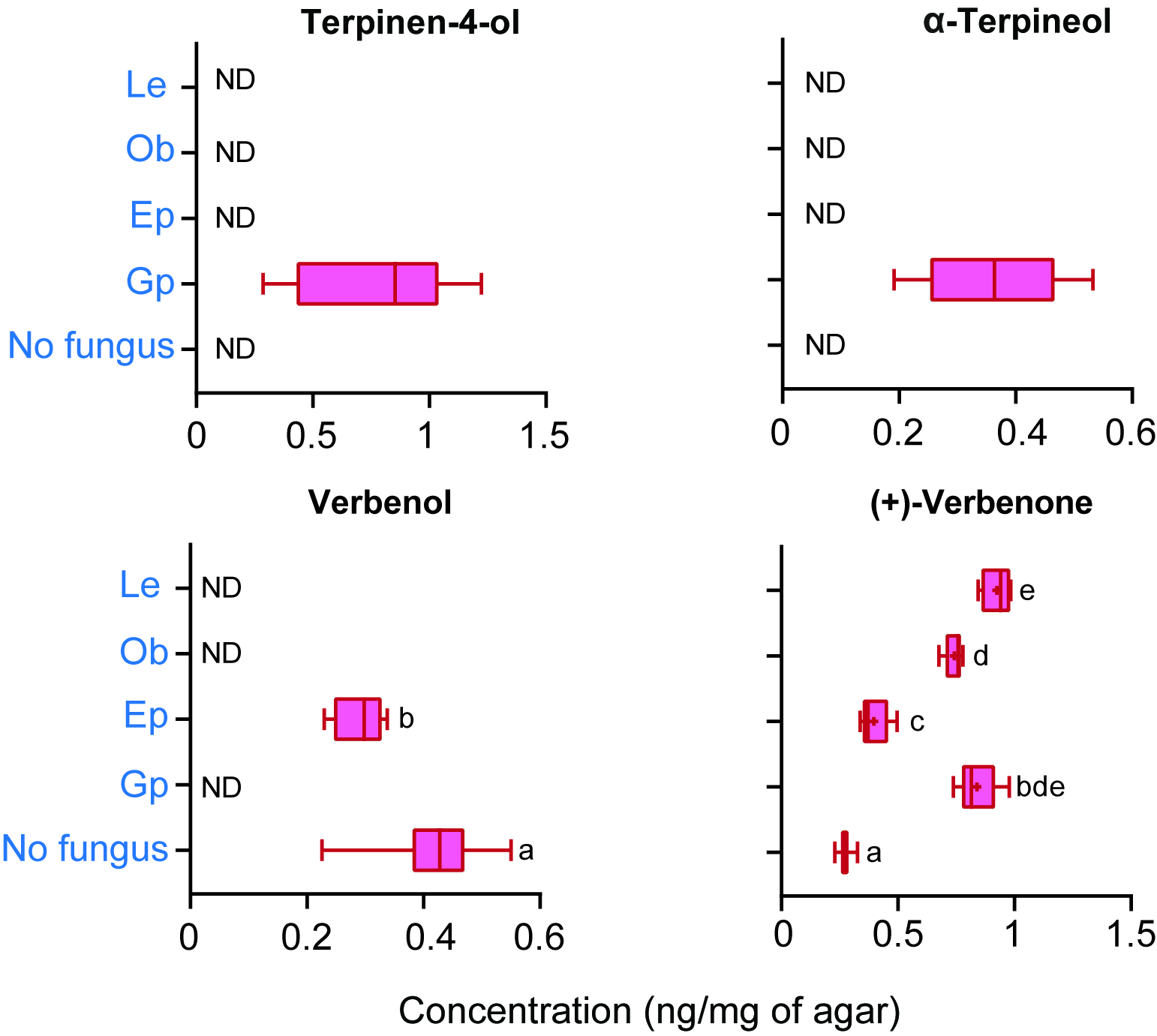

**Figure S7**: Metabolites of (+)-α-pinene produced by various *I. typographus* symbiotic fungi growing on potato dextrose agar after this monoterpene was administered to cultures of each species. Amounts of metabolites were determined after hexane extraction of the agar. Error bars represent SEM (*n* = 5). ND=not detected. Different lowercase letters denote significant differences between treatments (ANOVA, Sidak’s test; *P<*0.05). Fungal abbreviations: *E. polonica* (Ep), *L. europhioides* (Le), *G. penicillata* (Gp), *O. bicolor* (Ob).

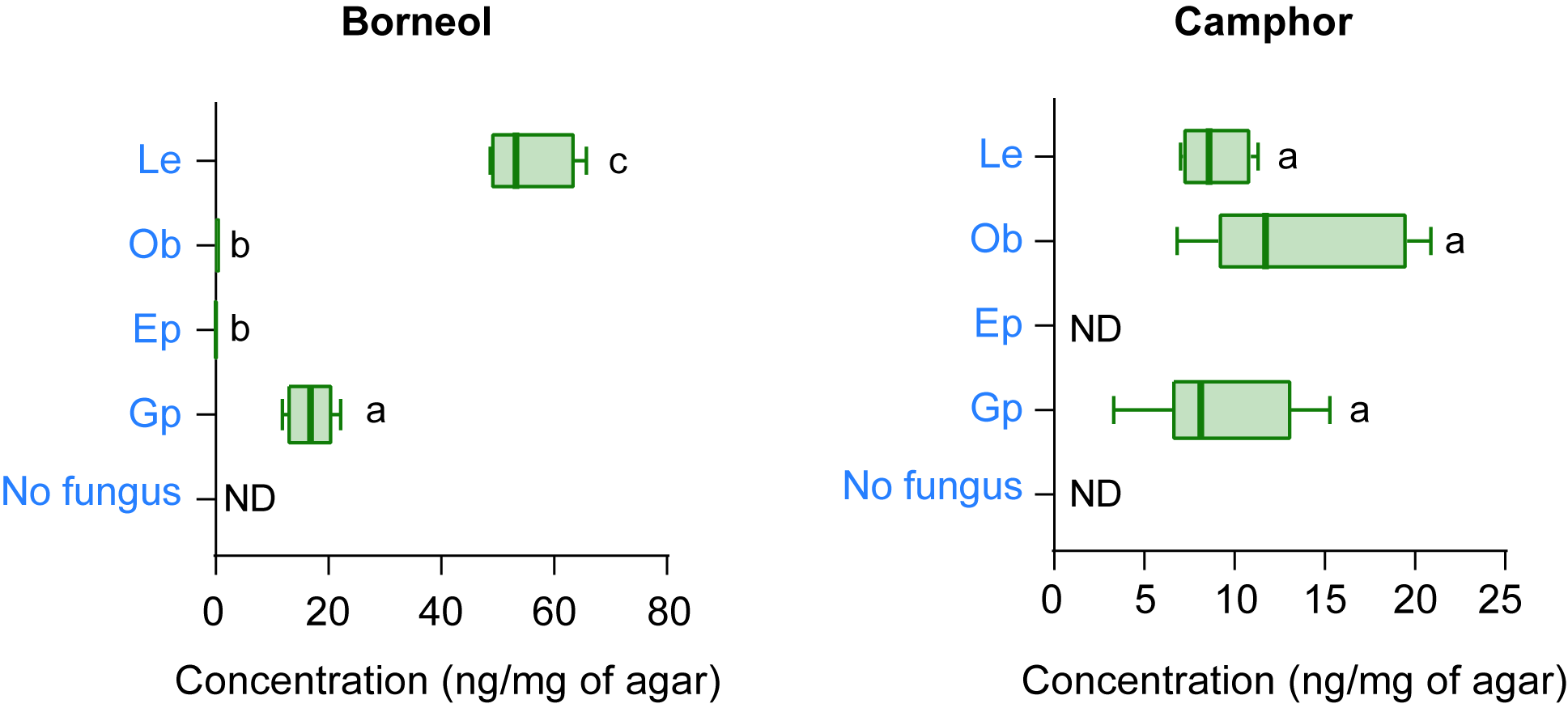

**Figure S8**: Metabolites of (-)-bornyl acetate produced by various *I. typographus* symbiotic fungi growing on potato dextrose agar after this monoterpene was administered to cultures of each species. Amounts of metabolites were determined after hexane extraction of the agar. Error bars represent SEM (*n* = 5). ND=not detected. Different lowercase letters denote significant differences between treatments (ANOVA, Sidak’s test; *P<*0.05). Fungal abbreviations: *E. polonica* (Ep), *L. europhioides* (Le), *G. penicillata* (Gp), *O. bicolor* (Ob).

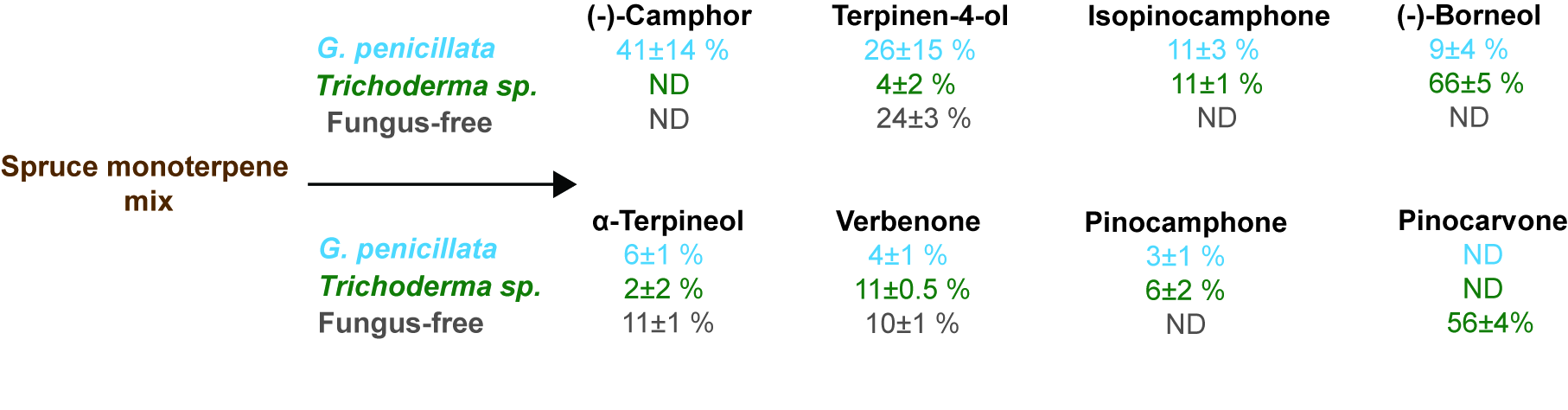

**Figure S9:** Relative proportion of oxygenated monoterpenes produced by the bark beetle symbiont *G. penicillata*, a saprophyte *Trichoderma sp.* and a fungus-free control potato dextrose agar medium amended with a mix of spruce monoterpenes (see Table Sx).

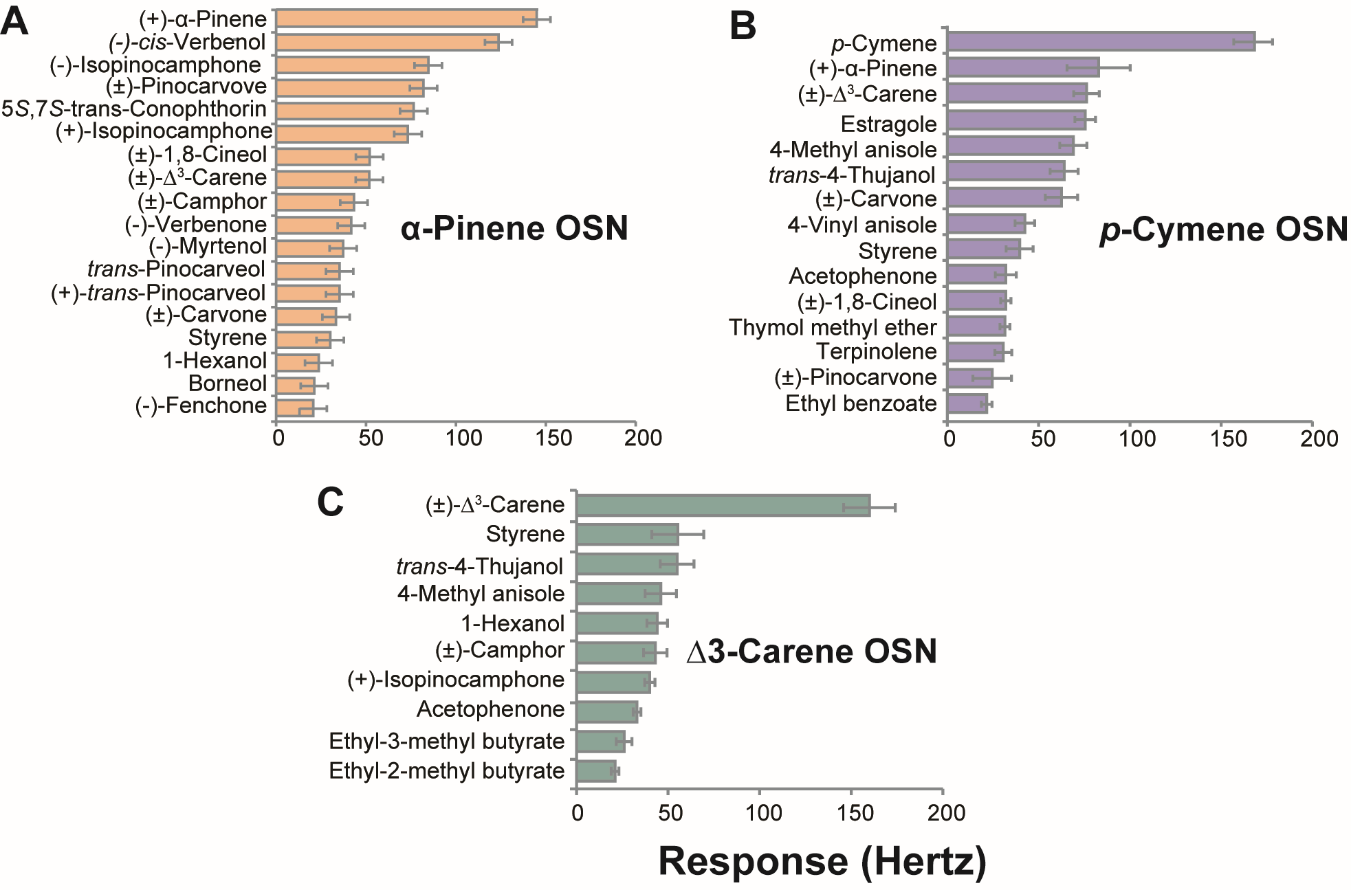

**Figure S10**: Response spectra of olfactory sensory neuron (OSN) classes (originally characterized in [59]) with primary responses to **(A)** (+)-α-pinene (*n* = 14), **(B)** *p*-cymene (*n* = 9), and **(C)** Δ3-carene (*n =* 4) to their respective most active ligands at the 10 µg screening dose (ligands eliciting responses <20 Hz are not shown). In addition to responses to the primary ligands, which are monoterpene hydrocarbons, these OSN classes show comparatively strong secondary responses to oxygenated monoterpenes produced by symbiotic fungi from host tree monoterpenes. Error bars represent SEM.

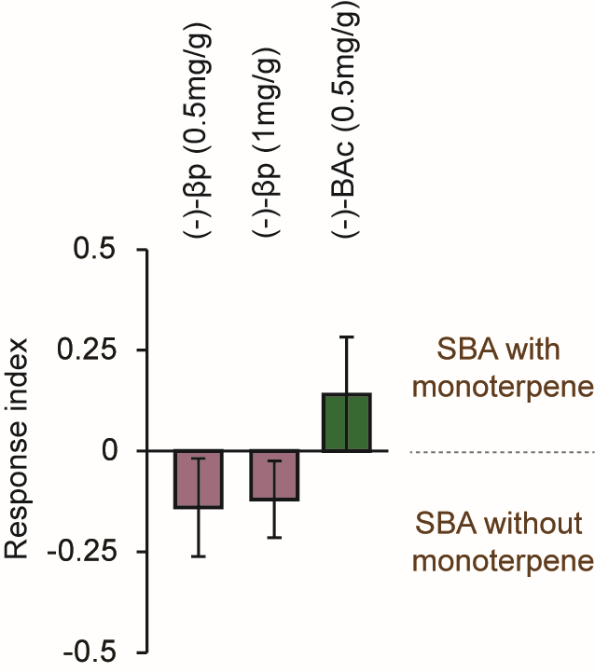

**Figure S11:** Adult beetles did not discriminate between spruce bark agar (SBA) enriched with monoterpenes and unenriched spruce agar. Error bars represent SEM (*n* = 25 for each trial)

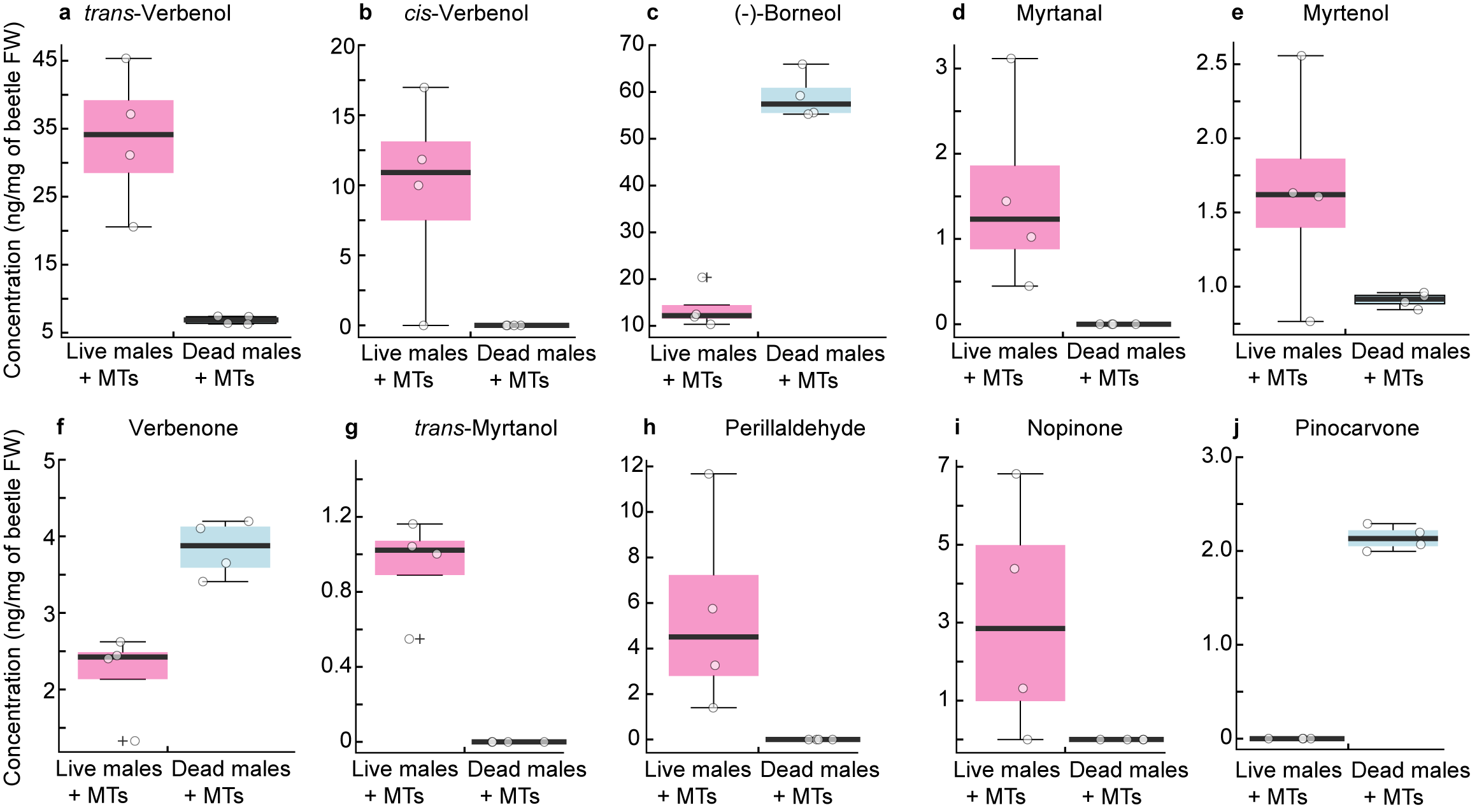

**Figure S12:** Concentration of the oxygenated monoterpenes (a) *trans*-verbenol, (b) *cis*-verbenol, (c) borneol, (d) myrtanal, (e) myrtenol, (f) verbenone, (g) *trans*-myrtanol, (h) perillaldehyde, (i) nopinone, and (j) pinocarvone produced by live and dead male *I. typographus* fumigated with the mix of spruce monoterpenes listed in Table S8.

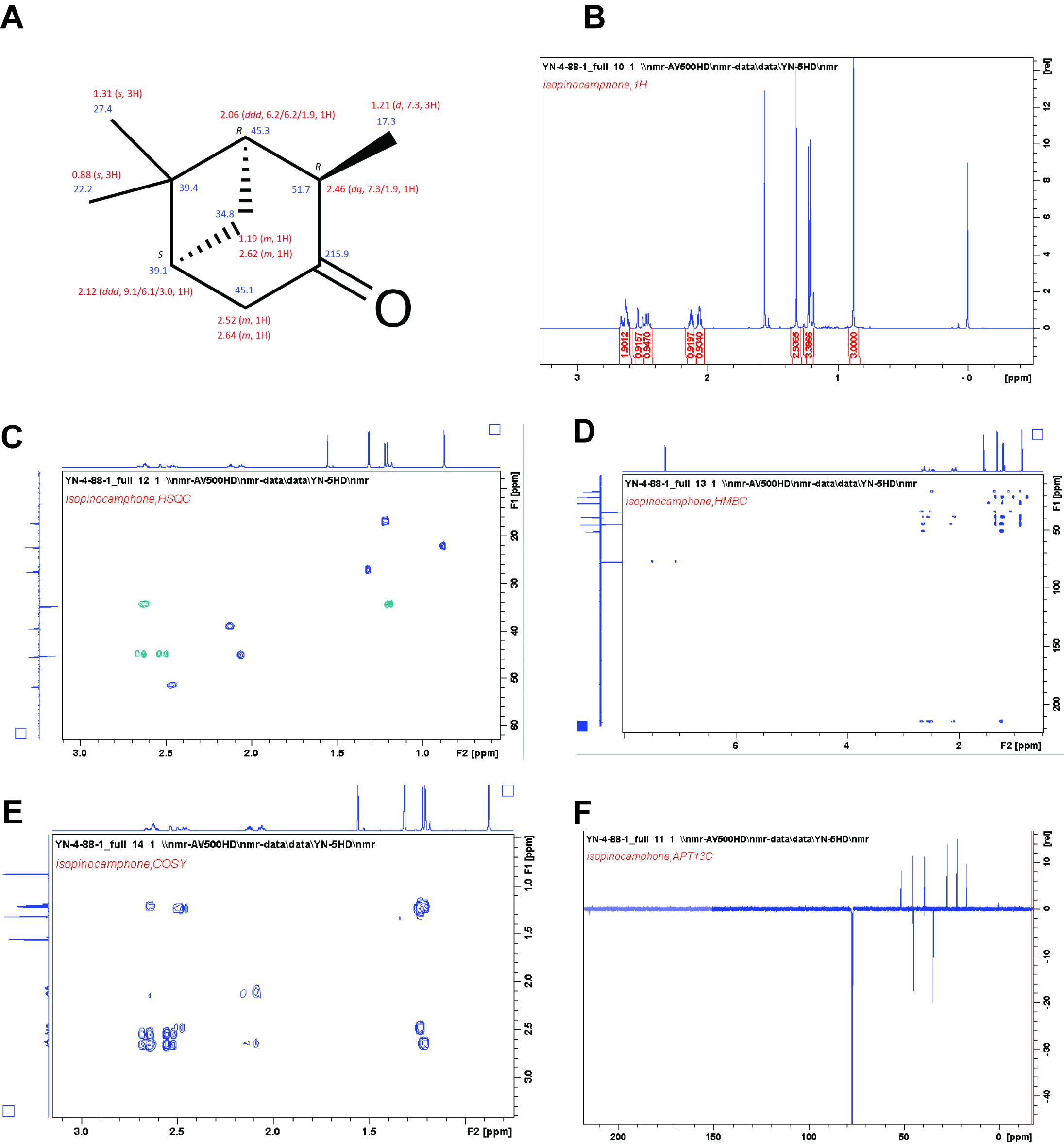

**Figure S13**: Confirmation of the structure of synthesized (+)-isopinocamphone by NMR. **(A)** ^1^H and ^13^C signal assignments in deuterated chloroform (CDCl_3_), **(B)** ^1^H NMR spectrum in CDCl_3_, **(C)** Phase sensitive heteronuclear single quantum coherence (HSQC) in CDCl_3_, **(D)** Heteronuclear multiple bond correlation (HMBC) in CDCl_3_, **(E)** Correlated spectroscopy (COSY) in CDCl_3_, **(F)** ^13^C attached proton test (APT) in CDCl_3_

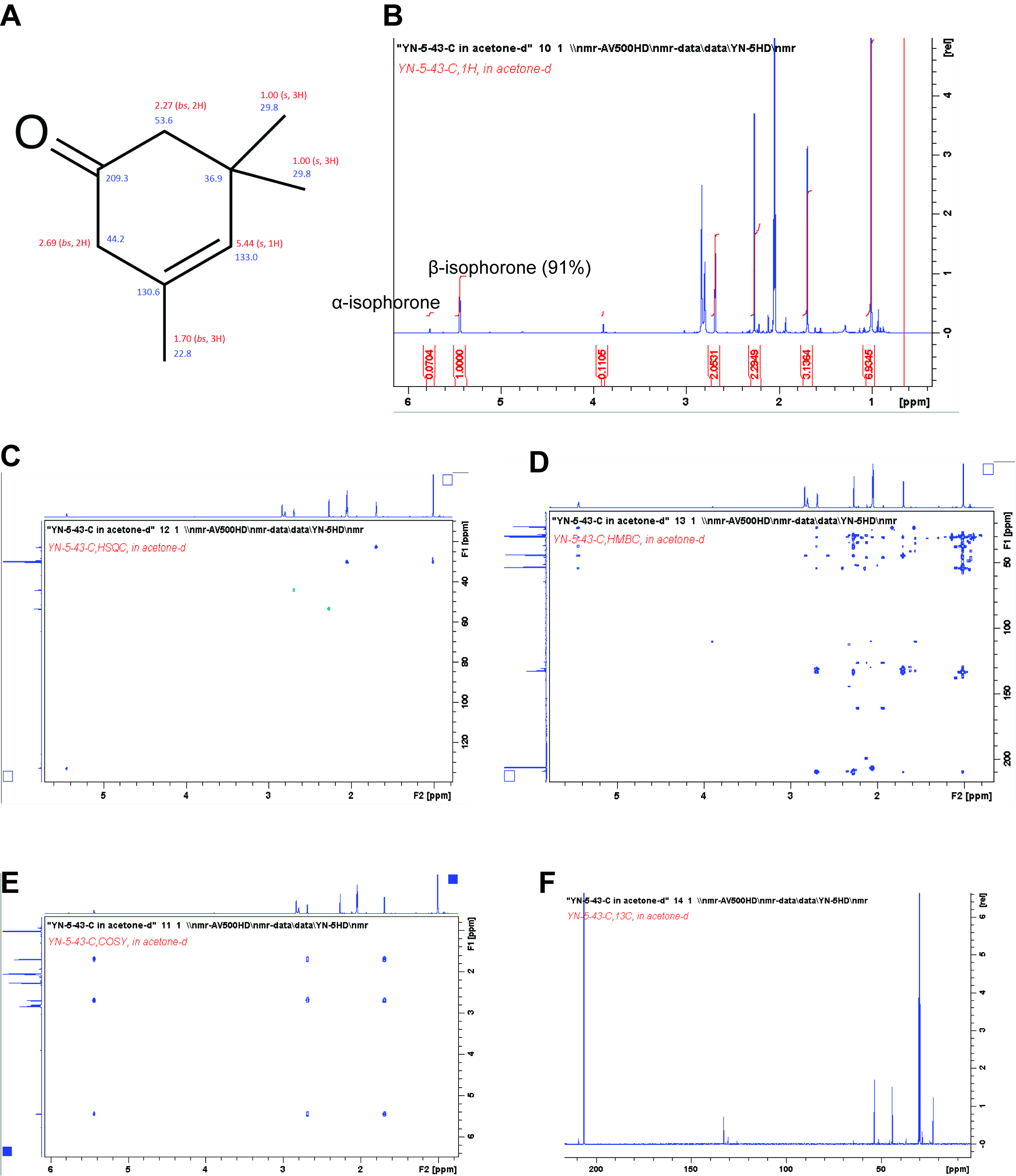

**Figure S14:** Confirmation of the structure of synthesized β-isophorone by NMR. **(A)** ^1^H and ^13^C signal assignments in acetone-*d_6_*, **(B)** ^1^H NMR spectrum in acetone-*d_6_*, purity- 91 % **(C)** Phase sensitive heteronuclear single quantum coherence (HSQC) in acetone-*d_6_*, **(D)** Heteronuclear multiple bond correlation (HMBC) in acetone-*d_6_*, **(E)** Correlated spectroscopy (COSY) in acetone-*d_6_*, **(F)** ^13^C spectrum in acetone-*d_6_*
