## Supplemental table for "Bark beetles locate fungal symbionts by detecting volatile fungal metabolites of host tree resin monoterpenes"

| ***Table S1***: Purity, source and biological origins of chemicals used in the experiments | | | |
| --- | --- | --- | --- |
| **Chemical** | **Purity (%)** | **Chemical source^1^** | **Biological origin** |
| Acetoin | >99 | Supelco | Fungi* |
| Acetophenone | 99 | Acros | Beetle/fungi^#^ |
| Amitinol | 96 | 1 | Beetle |
| Anisole | >99 | Sigma-Aldrich | Fungi^#^ |
| Benzaldehyde | >99 | Merck | Non-host/Fungi^*,#^ |
| Benzyl acetate | >99 | Aldrich | Fungi^*^ |
| Benzyl alcohol | >99 | Aldrich | Non-host/Fungi^*, #^ |
| Borneol | 94 | 2 | Fungi^#^ |
| (±)-2,3-Butanediol | >99 | Sigma-Aldrich | Fungi^*^ |
| (±)-Camphor | >99 | Aldrich | Host/fungi^#^ |
| (±)-∆3-Carene | 93 | Aldrich | Host |
| (±)-Carvone | >99 | Fluka | Fungi^#^ |
| (*E*)-β-Caryophyllene | >99 | Sigma | Host/fungi^*^ |
| β-Caryophyllene oxide | >99 | Givaudan-Roure | Fungi^*^ |
| Chavicol | >99 | Givaudan-Roure | Fungi^#^ |
| (±)-1,8-Cineole | >99 | Aldrich | Host |
| Cinnamaldehyde | >99 | Aldrich | Fungi^#^ |
| Cinnamyl alcohol | >99 | Aldrich | Fungi^#^ |
| Citral (*E* & *Z* mix) | >93 | Fluka | Fungi^*^ |
| (-)-Citronellol | >99 | Aldrich | Fungi^*^ |
| (+)-Citronellol | >99 | Aldrich | Fungi^*^ |
| β-Citronellyl acetate | >83 | 2 | Fungi^*^ |
| 5*S*,7*S*-*trans*-Conophthorin | 92 | 1 | Non-host/fungi^*,#^ |
| Coumaran | >99 | Sigma-Aldrich | Fungi^#^ |
| *p-*Cymene | >99 | Acros | Host |
| 4-Decanolide | ≥96 | Aldrich | Fungi^*^ |
| (-)-2,3-Dihydrofarnesol | >96 | 2 | Fungi^*^ |
| 3,4-Dimethoxytoulene | >98 | Sigma-Aldrich | Host/Fungi^#^ |
| Estragole | >99 | Fluka | Host/Fungi^#^ |
| Ethyl acetate | >99 | Sigma | Fungi^*^ |
| Ethyl benzoate | >99 | Fluka | Fungi^*^ |
| Ethyl butanoate | >99 | Firmenich | Fungi^*^ |
| Ethyl cinnamate | >99 | Aldrich | Fungi^#^ |
| 4-Ethyl guaiacol | >98 | Sigma | Fungi^#^ |
| Ethyl isobutyrate | >99 | Sigma-Aldrich | Fungi^*^ |
| Ethyl propionate | >99 | Aldrich | Fungi^*^ |
| Ethyl-2-methylbutyrate | >99 | Sigma-Aldrich | Fungi^*^ |
| Ethyl-3-methylbutyare | >99 | Sigma-Aldrich | Fungi^*^ |
| *p-*Eugenol | >99 | Fluka | Fungi^#^ |
| (*E,E*)-α-Farnesene | 95 | Fluka | Host/Fungi^*^ |
| (*E,E*)-Farnesol | 95 | Fluka | Fungi^*^ |
| (-)-Fenchol | >99 | 2 | Fungi^#^ |
| (-)-Fenchone | >99 | Fluka | Fungi^#^ |
| Geranyl acetate | >90 | Fluka | Fungi^*^ |
| Geranylacetone | >97 | Fluka | Non-host/fungi^*^ |
| 1-Hexanol | >99 | Fluka | Non-host/fungi^*^ |
| (±)-Ipsdienol | 94 | Bedoukian | Beetle |
| (±)-Ipsenol | 97 | Bedoukian | Beetle |
| Isoamyl acetate | >98 | Aldrich | Fungi^*^ |
| Isobutyl acetate | >99 | Fluka | Fungi^*^ |
| Isoeugenol | >99 | Aldrich | Fungi^#^ |
| α-Isophorone | >99 | Acros | NA |
| β-Isophorone | >90 | 3 | Beetle |
| (-)-Isopinocamphone | 96 | 4 | Host/fungi^#^ |
| (+)-Isopinocamphone | 92 | 3,4 | Host/fungi^#^ |
| 4-Methyl anisole | >99 | Aldrich | Fungi^#^ |
| Methyl cinnamate | >99 | Sigma-Aldrich | Fungi^#^ |
| Methyl eugenol | >90 | Sigma | Host/Fungi^#^ |
| 2-Methyl-1-butanol | >99 | Aldrich | Fungi^*^ |
| 3-Methyl-1-butanol | 99 | Aldrich | Fungi^*^ |
| 2-Methyl-3-buten-2-ol | >99 | Acros | Beetle/fungi^*^ |
| Myrcene | 95 | Fluka | Host |
| (-)-Myrtenol | 99 | 2 | Beetle/fungi^#^ |
| Neryl acetate | ≥97 | Aldrich | Fungi^*^ |
| (±)-3-Octanol | >99 | Acros | Non-host/fungi^*^ |
| (±)-Octen-3-ol | 98 | Acros | Non-host/fungi^*^ |
| Phenol | >99 | Sigma-Aldrich | Fungi^#^ |
| 3-Phenyl propanol | >99 | Sigma | Fungi^*^ |
| 2-Phenylethanol | >95 | Merck | Beetle/Fungi^*^ |
| 2-Phenylethyl acetate | >99 | Aldrich | Fungi^*^ |
| N-Phenylformamide | >99 | Alfa Aesar | Fungi^*^ |
| (+)-α-Pinene | >99 | Janssens chimica | Host |
| (*E*)-Pinocarveol | 67 | 2 | Fungi^#^ |
| (+)-(*E*)-Pinocarveol | 99 | 1 | Fungi^#^ |
| (±)-Pinocarvone | ~51^$^ | Synergy Corp. | Beetle/fungi^#^ |
| Propyl propionate | >99 | Acros | Fungi^*^ |
| Salicylaldehyde | >99 | Aldrich | Fungi^#^ |
| Styrene | 99 | Fluka | Fungi^#^ |
| Terpinolene | 97 | Fluka | Host |
| (+)-*trans*-4-Thujanol | >99 | Acros | Host/ Fungi^#^ |
| (-)-Terpinen-4-ol | >99 | Acros | Fungi^#^ |
| (+)-Terpinen-4-ol | >99 | Acros | Fungi^#^ |
| (-)-α-Terpineol | >99 | Fluka | Host/ Fungi^#^ |
| (+)-α-Terpineol | >99 | Fluka | Host/ Fungi^#^ |
| Thymol methyl ether | >99 | Sigma-Aldrich | Host |
| Toluene | >99 | Aldrich | Fungi^#^ |
| Vanillin | >99 | Sigma | Fungi^#^ |
| (-)-*cis*-Verbenol | 95 | Borregaard | Beetle |
| (+)-*trans*-Verbenol | 92 | SCM | Beetle |
| (-)-*trans*-Verbenol | 97 | Sci Tech | Beetle |
| (-)-Verbenone | >99 | Fluka | Beetle/fungi^#^ |
| 4-Vinyl anisole | >99 | Sigma-Aldrich | Fungi^#^ |
| ^1^Code to numbers: (1) Gift of Wittko Francke, University of Hamburg, Hamburg, Germany, (2) gift of Gunnar Bergström, University of Göteborg, Göteborg, Sweden, (3) synthesized in this study (4) gift of Rikard Unelius, Linnaeus University, Kalmar, Sweden. | | | |
| ^#^Fungal metabolite of host tree volatile | |  |  |
| ^*^Fungal volatile produced *de novo* | |  |  |
| ^$^None of the other components were active | |  |  |
| NA= Not applicable |  |  |  |

***Table S2*:** Emission of volatile organic compounds identified from the headspace collection of fresh spruce bark four days after inoculation with different fungi. Analyses were conducted using GC-FID. Compounds with significant *P* values are highlighted in **bold.**

|  | **RT^#^** | ***F^$^*** | ***P^$^*** | **Emission rate at 4 days post-inoculation (pg mg dry weight of bark^-1^ h^-1^)** | | | | | |
| --- | --- | --- | --- | --- | --- | --- | --- | --- | --- |
|  |  |  |  | **Uninfected** | ***E. polonica*** | ***G. penicillata*** | ***L. europhioides*** | ***O. bicolor*** | ***O. piceae*** |
| ***Monoterpenes*** |  |  |  |  |  |  |  |  |  |
| **Santene** | 5.70 | 7.97 | **<0.001** | 2.91±0.32^bc^ | 2.59±0.14^c^ | 2.75±0.22^bc^ | 4.66±0.51^ab^ | 5.41±0.7^a^ | 5.81±0.92^a^ |
| Tricyclene | 6.53 | 2.46 | 0.073 | 7.22±1.49 | 8.92±1.64 | 8.89±0.54 | 20.08±8.49 | 15.1±2.26 | 19.11±2.9 |
| **α-Thujene** | 6.68 | 4.25 | **0.01** | 4.8±0.86^b^ | 6.4±1.04^ab^ | 8.42±1.06^ab^ | 17.1±6.1^ab^ | 12.32±1.52^ab^ | 14.16±2^a^ |
| **α-Pinene** | 6.81 | 3.94 | **0.014** | 1308±236^b^ | 1942±424^ab^ | 2036±211^ab^ | 3733±1029^ab^ | 3471±531^ab^ | 4059±634^a^ |
| Camphene | 7.15 | 2.13 | 0.109 | 60.04±15.24 | 73.63±16.8 | 62.09±6.11 | 131±51.59 | 123±19.32 | 160±26.84 |
| ***o*-Cymene** | 7.70 | 3.3 | **0.027** | 2.87±1.53^a^ | 2.55±0.69^b^ | 7.76±1.61^ab^ | 5.61±1.42^ab^ | 5.08±0.8^ab^ | 7.55±1.37^ab^ |
| Sabinene | 7.76 | 2.26 | 0.134 | 20.63±5.01 | 33.89±5.82 | 24.28±3.88 | 13.02±7.63 | ND | ND |
| **β-Pinene** | 7.83 | 4.36 | **0.009** | 1871±313^b^ | 2885±566^ab^ | 3180±586^ab^ | 4696±681^ab^ | 5427±706^a^ | 5711±855^a^ |
| **Myrcene** | 8.20 | 7.57 | **<0.001** | 45.1±4.6^c^ | 70.94±12.72^bc^ | 99.5±30.6^abc^ | 189±27.84^a^ | 181±28^a^ | 150±17.58^ab^ |
| **α-Phellandrene** | 8.50 | 14.74 | **<0.001** | 4.36±0.46^c^ | 4.78±0.25^c^ | 7±0.75^bc^ | 12.53±1.58^a^ | 10.15±0.83^ab^ | 7.92±0.59^ab^ |
| Δ3-Carene | 8.64 | 1.89 | 0.146 | 83.99±20.9 | 116±24.87 | 182±23.94 | 211±48.86 | 114±11.09 | 155±15.23 |
| **α-Terpinene** | 8.80 | 2.97 | **0.042** | 3.24±0.21^b^ | 3.74±0.63^ab^ | 3.66±1.51^ab^ | 10.84±3.25^a^ | 4.84±0.54^ab^ | 6.09±0.96^ab^ |
| ***p*-Cymene** | 9.00 | 4.45 | **0.008** | 32.3±7.18^c^ | 47.09±9.09^bc^ | 66.61±13.95^abc^ | 73.87±9.06a^bc^ | 91.71±9.27^ab^ | 111±18.52^a^ |
| **β-Phellandrene** | 9.10 | 3.59 | **0.02** | 501±77.5^b^ | 773±161^ab^ | 726±159^ab^ | 1390±255^ab^ | 1395±196^a^ | 1161±152^ab^ |
| **γ-Terpinene** | 9.84 | 3.08 | **0.035** | 3.76±0.39^b^ | 3.93±1.23^ab^ | 4.04±0.98^ab^ | 8.95±1.41^a^ | 6.03±0.82^ab^ | 6.17±0.83^ab^ |
| ***Spiroketals/ others*** |  |  |  |  |  |  |  |  |  |
| ***endo*-1,3-Dimethyl-2,9-dioxabicyclo[3.3.1]nonane** | 9.36 | 8.16 | **<0.001** | ND | 1.85±1.17^c^ | 145±41.82^a^ | 82.7±10.5^abc^ | 126±16.49^ab^ | 34.67±13.58^bc^ |
| ***exo*-1,3-Dimethyl-2,9-dioxabicyclo[3.3.1]nonane** | 10.77 | 12.22 | **<0.001** | ND | 4.11±1.15^c^ | 144±32.23^ab^ | 203±28.94^a^ | 241±33.79^a^ | 72.88±20.94^bc^ |
| Nonanal | 10.95 | 0.55 | 0.738 | 8.99±1.07 | 7.12±0.62 | 8.19±0.47 | 9.33±0.75 | 8.95±1.23 | 9.07±1.31 |
| ***Oxygenated monoterpenes*** |  |  |  |  |  |  |  |  |  |
| ***trans*-4-Thujanol** | 10.06 | 4.93 | **0.005** | 1.85±0.35^b^ | 2.43±0.49^b^ | 7.69±2.85^ab^ | 11.61±1.43^a^ | 9.23±1.48^ab^ | 9.62±1.55^ab^ |
| **Linalool oxide** | 10.19 | 8.05 | **<0.001** | 2.66±0.36^c^ | 3.9±0.62^bc^ | 10.53±1.41^abc^ | 20.19±4.05^a^ | 22.98±3.45^a^ | 16.58±3.39^ab^ |
| **Terpinolene/Fenchone/Cymenene^1^** | 10.57 | 7.97 | **<0.001** | 12.88±2.68^d^ | 17.49±2.31^cd^ | 24.6±3.34^bcd^ | 54.18±10.15^a^ | 47.3±5.36^ab^ | 40.2±4.69^abc^ |
| **Fenchol** | 11.17 | 13.78 | **<0.001** | 4.32±1.52^cd^ | 0.58±0.38^d^ | 5.77±1.04^bcd^ | 9.86±1.42^bc^ | 14.38±1.85^ab^ | 20.73±3.02^a^ |
| **β-Thujone** | 11.26 | 8.81 | **<0.001** | 0.41±0.35^c^ | 1.72±0.22^bc^ | 2.79±0.23^bc^ | 2.45±0.83^bc^ | 7.21±1.34^a^ | 4.45±0.47^ab^ |
| Unknown monoterpene #1 | 11.48 | 1.55 | 0.224 | 0.64±0.33 | 1.25±0.38 | 83.05±18.84 | 19.92±2.83 | 4.5±0.61 | 37.48±15.23 |
| ***trans*-Pinocarveol** | 11.78 | 10.75 | **<0.001** | 22.38±5.09^d^ | 48.71±7.24^cd^ | 83.4±9.73^bcd^ | 96.03±15.7^abc^ | 182.7±24.65^a^ | 144.18±27.28^ab^ |
| **Camphor** | 11.92 | 77.48 | **<0.001** | 26.32±3.4^c^ | 96.46±18.78^c^ | 1280±140^b^ | 1565±249^ab^ | 2537±256^a^ | 859±98.63^b^ |
| **Isoborneol** | 12.22 | 8.94 | **0.002** | ND | ND | 9.94±1.06^b^ | 11±1.37^b^ | 13.7±1.51^b^ | 30.24±6.16^a^ |
| **Pinocamphone** | 12.30 | 5.85 | **0.002** | 93.16±16.52^b^ | 147±24.64^ab^ | 165±7.67^ab^ | 260±53.62^a^ | 281±28^a^ | 287±35.54^a^ |
| Pinocarvone | 12.36 | 0.62 | 0.46 | ND | ND | ND | ND | 55.9±5.39 | 67.82±11.1 |
| ***endo*-Borneol** | 12.43 | 36.28 | **<0.001** | 13.15±4.94^d^ | 80.66±12.15^cd^ | 225±27.93^bc^ | 375±50.6^b^ | 175±30.35^bcd^ | 166±5396.75^a^ |
| **Isopinocamphone** | 12.63 | 14.57 | **<0.001** | 41.9±9.38^d^ | 88.28±22.45^cd^ | 137±7.53^cd^ | 453±100.9^a^ | 197±21.96^bcd^ | 368±52.31^ab^ |
| **Terpinen-4-ol** | 12.70 | 10.72 | **<0.001** | ND | 7.09±1.87^c^ | 94.91±55.52^ab^ | 22.17±4.43^bc^ | 101±17.52^ab^ | 136±39.66^a^ |
| *p*-Cymene-8-ol | 12.88 | 2.08 | 0.115 | 3.32±1.18 | 7.56±1.46 | 23.89±7.32 | 43.53±7.31 | 55.52±11.76 | 31.03±10.76 |
| α-Terpineol | 12.95 | 1.65 | 0.23 | ND | ND | 5.2±1.09 | 16.2±3.45 | 13.14±4.4 | 12.85±2.78 |
| **Verbenone** | 13.18 | 6.37 | **<0.001** | 41.73±6.51^b^ | 62.7±4.64^b^ | 95.64±8.62^b^ | 126±18.92^ab^ | 216±34.26^a^ | 127±36.35^ab^ |
| **Thymol methyl ether** | 14.22 | 7.73 | **<0.001** | 3.12±0.63^c^ | 8.22±1.19^bc^ | 20.85±2.69^abc^ | 36.23±4.1^ab^ | 47.37±7.55^a^ | 44.02±12.19^a^ |
| ***p*-Menth-2-en-7-ol** | 14.55 | 5.78 | **0.005** | ND | 12.82±5.13^c^ | 32.37±5.65^abc^ | 52.91±5.65^ab^ | 59.88±12.28^a^ | 15.15±8.26^bc^ |
| **Myrtanol** | 14.58 | 15.66 | **<0.001** | 6.43±0.63^c^ | 10.1±1^c^ | 17.05±2.62^bc^ | 38.95±5.71^ab^ | 61.37±12.71^a^ | 72.98±19.9^a^ |
| **Bornyl acetate** | 15.20 | 15.11 | **<0.001** | 14.09±2.77^bc^ | 53.18±5.36^a^ | 44.41±6.59^a^ | 23.09±1.55^ab^ | 7.6±0.41^c^ | 51.85±11.15^a^ |
| Thymol | 15.30 | 0.99 | 0.453 | 0.54±0.27 | 5.87±1.24 | 13.97±3.84 | 45.19±6.78 | 10.41±2.14 | 13.99±4.86 |
| ***Sesquiterpenes*** |  |  |  |  |  |  |  |  |  |
| α-Longipinene | 16.63 | 2.15 | 0.106 | 10.62±1.17 | 14.7±1.98 | 13.65±3.15 | 28.01±7.48 | 21.66±2.12 | 17.53±3.11 |
| Unknown SqT #1 | 16.69 |  |  | ND | ND | ND | ND | ND | 11.98±2.39 |
| Cyclosativene | 17.08 | 2.21 | 0.099 | 6.05±0.93 | 8.48±0.82 | 7.27±1.41 | 12.64±2.86 | 10.39±0.89 | 6.42±0.97 |
| **α-Cubebene** | 17.17 | 2.99 | **0.039** | 2.65±0.36^a^ | 3.38±0.33^a^ | 2.73±0.6^a^ | 8.12±2.57^a^ | 6.4±0.53^a^ | 3.36±0.66^a^ |
| 6-Protoilludene | 17.26 |  |  | ND | ND | ND | ND | ND | 188.97±39.53 |
| Gurjunene | 17.48 | 2.23 | 0.096 | 5.48±1.25 | 6.27±0.8 | 4.09±1.37 | 10.09±1.28 | 7.64±1.25 | 8.72±1.57 |
| Isolongifolene | 17.81 | 2.28 | 0.09 | 39.35±5.05 | 59.72±8.29 | 49.94±9.95 | 86.56±20.9 | 71.7±6.33 | 40.86±7.04 |
| α-Cedrene | 17.95 | 2.75 | **0.051** | 3.57±0.59^ab^ | 5.1±0.65^ab^ | 4.68±0.71^ab^ | 8.35±2.16^a^ | 5.41±0.38^ab^ | 3.26±0.57^b^ |
| (*E*)-β-Caryophyllene | 18.10 | 1.87 | 0.151 | 5.56±0.9 | 7.83±0.93 | 9.22±1.19 | 16.3±4.94 | 10.42±0.83 | 11.29±2.17 |
| (*E*)-β-Caryophyllene (fungus) | 18.48 | 1.09 | 0.336 | ND | ND | 6.08±1.64 | 8.62±1.33 | ND | ND |
| Δ-Cadinene | 18.71 | 1.22 | 0.341 | 1.95±0.49 | 3.28±0.32 | 2.14±0.34 | 3.54±0.59 | 2.91±0.13 | 2.63±0.8 |
| **Humulene/(*E*)-β-Farnesene^1^** | 18.81 | 3.94 | **0.014** | 4.06±0.67^b^ | 5.38±0.46^ab^ | 7.99±0.97^ab^ | 9.49±1.31^a^ | 9.87±0.57^a^ | 6.36±1.58^ab^ |
| Amorphene | 19.25 | 2.34 | 0.084 | 9.48±1.23 | 12.4±1.42 | 10.5±1.35 | 25.18±9.14 | 21.88±1.79 | 11.47±2.73 |
| Cubenene | 19.35 | 1.68 | 0.19 | 22.25±1.77 | 34.05±3.73 | 33.1±7.12 | 75.84±30.03 | 49.04±5.76 | 31.45±8.37 |
| α-Muurolene | 19.72 | 1.72 | 0.18 | 10.17±1.69 | 15.65±2.01 | 11.35±1.17 | 22.48±7.08 | 17.14±1.27 | 9.74±2.5 |
| **Unknown SqT #2** | 20.09 | 3.33 | **0.027** | 1.8±0.27^ab^ | 1.53±0.11^ab^ | 1.54±0.04^ab^ | 2.3±0.15^a^ | 2.2±0.21^ab^ | 1.14±0.34^b^ |
| Δ-Cadinene | 20.17 | 2.7 | 0.054 | 30.38±2.83 | 53.94±7.58 | 42.88±4.92 | 88.63±23.85 | 80.15±6.76 | 50.83±14.2 |
| **Germacrene B** | 20.84 | 3.07 | **0.035** | 2.91±0.58^a^ | 5.81±0.82^a^ | 5.52±0.85^a^ | 5.24±0.69^a^ | 3.19±0.43^a^ | 2.67±0.81^a^ |
| Caryophyllene oxide | 21.94 | 4 | 0.093 | ND | ND | 24.06±6.27 | 9.16±1.76 | ND | ND |

^#^- Estimated retention time using the method developed for GC-FID described in the materials and methods section of this manuscript

***^$^*** Significant differences between species are denoted by small letters (ANOVA, followed by Tukey’s test, *P<0.05)*

^1^-Compounds co-eluted from the GC column.

Unknown monoterpene #1: M^+^ = *m/z* 148

Unknown SqT (sesquiterpene) #1: M^+^ = *m/z* 204.2

Unknown SqT (sesquiterpene) #2: M^+^ = *m/z* 204

***Table S3***. Relative amounts (mean ± SE, n=5) of volatiles from uninfected bark detected after various time periods (4, 8, 12 and 18 days) from the beginning of an experiment with fungal inoculation. Data from the control uninfected treatment are presented here. Data for fungal treatments are given in Tables S3-S6. Volatiles were collected on polydimethylsiloxane tubes for 2 hours and were subjected to GC-MS analysis (see materials and methods section for details). ND=not detected, NA=not analyzed, TR= trace amounts (<500 TIC counts).

| ***Compounds*** | **RT^#^** | ***F^$^*** | ***P^$^*** | **Uninfected bark peak area (*10^4^ TIC counts)** | | | |
| --- | --- | --- | --- | --- | --- | --- | --- |
|  |  |  |  | **4d** | **8d** | **12d** | **18d** |
| ***Aliphatics*** |  |  |  |  |  |  |  |
| 2-Butanone | 1.85 | - | - | ND | ND | ND | 2.86±0 |
| 2-Methyl-3-buten-2-ol | 1.93 | - | - | ND | ND | ND | 0.08±0 |
| Ethyl acetate | 1.95 | - | - | ND | ND | ND | ND |
| Isobutanol | 2.40 | - | - | ND | ND | ND | ND |
| Isopropyl acetate | 2.33 | - | - | ND | ND | ND | ND |
| **Acetoin** | 2.85 | 19.32 | **0.005** | 6.57±0.11(a) | 6.49±1.05(a) | 4.27±0.8(ab) | 0.09±0.05(b) |
| Ethyl propanoate | 2.88 | - | - | ND | 1.6±0.15 | ND | ND |
| 3-Methyl-1-butanol | 3.24 | - | - | ND | ND | 0.62±0.11 | 0.72 |
| Ethyl isobutyrate | 3.69 | - | - | ND | 0.07±0 | ND | ND |
| Isobutyl acetate | 3.99 | 1.16 | 0.341 | ND | 0.08±0.01 | 0.1±0.01 | ND |
| 2,3-Butanediol | 4.17 | - | - | ND | ND | ND | ND |
| Ethyl butanoate | 4.55 | - | - | ND | ND | 0.05±0 | ND |
| Ethyl but-2-enoate | 5.60 | - | - | ND | ND | ND | ND |
| Ethyl 2-methylbutyrate | 5.75 | - | - | ND | ND | ND | ND |
| 1-Hexanol | 6.25 | - | - | ND | ND | 0.14±0 | 0.07±0 |
| 3-Methyl-1-butyl acetate | 6.46 | - | - | ND | 10.78±0.54 | 0.04±0 | ND |
| Isopentyl-2-methylbutanoate | 12.47 | - | - | ND | ND | ND | ND |
| Isoamyl valerate | 12.60 | - | - | ND | ND | ND | TR |
| ***Aromatics*** |  |  |  |  |  |  |  |
| 2-Phenylethyl alcohol | 12.79 | - | - | ND | ND | ND | ND |
| 2-Phenylethyl acetate | 16.39 | - | - | ND | 0.07±0 | ND | ND |
| Citronellyl acetate | 18.58 | - | - | ND | ND | ND | ND |
| ***Spiroketals*** |  |  |  |  |  |  |  |
| *endo-*1,3-dimethyl-2,9-dioxabicyclo[3.3.1]nonane | 10.81 | - | - | ND | ND | ND | ND |
| *trans*-Conophthorin | 11.29 | - | - | ND | ND | ND | 0.03±0 |
| Brevicomin | 11.64 | - | - | ND | ND | ND | ND |
| *exo-*1,3-dimethyl-2,9-dioxabicyclo[3.3.1]nonane | 12.37 | - | - | ND | ND | 0.06±0.01 | 0.14 |
| ***Monoterpenes*** |  |  |  |  |  |  |  |
| Santene | 6.61 | 0.94 | 0.357 | 0.34±0.11 | 0.09±0.03 | 0.09±0.03 | 0.14±0 |
| **Tricyclene** | 7.67 | 6.94 | **0.027** | 2.57±0.35(a) | 0.58±0.14(ab) | 0.23±0.07(ab) | 0.41±0.11(b) |
| α-Thujene | 7.76 | 3.36 | 0.1 | 0.73±0.09 | 0.12±0.03 | 0.04±0.01 | 0.19±0.06 |
| **α-Pinene** | 7.94 | 8.77 | **0.016** | 465±89.61(a) | 122±29.58(ab) | 57.75±14.54(ab) | 72.2±14.92(b) |
| **Camphene** | 8.34 | 9.33 | **0.014** | 7.67±1.38(a) | 2.4±0.5(ab) | 1.06±0.23(ab) | 1.44±0.1(b) |
| **Verbenene** | 8.51 | 12.18 | **0.007** | 0.35±0.08(a) | 0.08±0.02(ab) | 0.05±0.01(b) | 0.04±0(b) |
| Sabinene | 9.50 | - | - | ND | ND | ND | ND |
| **β-Pinene** | 9.13 | 17.75 | **0.002** | 866±148(a) | 188±51.19(ab) | 76.37±25.05(b) | 68.05±10.8(b) |
| **β-Myrcene** | 9.54 | 6.44 | **0.035** | 7.48±1.31(a) | 1.62±0.63(a) | 1.05±0.36(a) | 1.08±0.04(a) |
| **α-Phellandrene** | 9.88 | 10.6 | **0.01** | 0.61±0.1(a) | 0.27±0.09(ab) | 0.03±0.03(ab) | 0.18±0(b) |
| α-Terpinene | 10.21 | - | - | 0.1±0 | ND | ND | ND |
| *p*-Cymene | 10.43 | 4.88 | 0.054 | 10.29±3.01 | 4.27±1.4 | 2.56±0.73 | 2.7±0.85 |
| **Limonene** | 10.51 | 7.9 | **0.02** | 26.08±5.15(a) | 6.91±1.85(ab) | 3.11±0.88(b) | 4.11±0.69(ab) |
| **β-Phellandrene** | 10.55 | 6.1 | **0.036** | 56.99±9.75(a) | 18.3±5.48(ab) | 7.48±2.11(ab) | 13.6±0.94(b) |
| γ-Terpinene | 11.37 | - | - | ND | ND | ND | ND |
| α-Terpinolene | 12.16 | - | - | ND | ND | ND | 0.09±0 |
| *p*-Cymenene | 12.19 | 0 | 0.983 | 0.1±0.02 | 0.06±0.02 | 0.1±0 | 0.09±0.05 |
| ***Oxygenated monoterpenes*** |  |  |  |  |  |  |  |
| **1,8-Cineole** | 10.61 | 10.71 | **0.011** | 6.27±1.55(a) | 2.8±0.94(ab) | 0.58±0.22(ab) | 0.02±0(b) |
| Linalool oxide | 11.73 | - | - | ND | ND | ND | TR |
| Fenchone | 12.15 | 0 | 0.961 | 0.54±0.28 | 0.68±0.27 | 0.39±0.19 | 0.59±0.15 |
| *trans*-4-Thujanol | 12.42 | - | - | ND | ND | ND | ND |
| *exo*-Fenchol | 12.82 | 0.03 | 0.875 | 0.09±0.03 | 0.06±0.02 | 0.05±0.02 | 0.09±0.05 |
| β-Thujone | 12.93 | - | - | 0.09±0.01 | TR | ND | ND |
| *p*-Isopropylcyclohexanol | 13.41 | - | - | ND | ND | ND | ND |
| *trans*-Pinocarveol | 13.48 | 1.36 | 0.274 | 0.27±0 | 0.08±0.01 | 0.27±0.03 | 0.18±0.05 |
| Camphor | 13.63 | 1.07 | 0.329 | 1.34±1.02 | 1.34±0.98 | 1.09±0.74 | 2.45±0.28 |
| Camphene hydrate | 13.73 | 0.49 | 0.71 | 0.22±0.06 | 0.2±0.06 | 0.16±0.05 | 0.08±0.01 |
| Pinocamphone | 14.43 | 0.46 | 0.513 | 2.71±0.87 | 1.43±0.7 | 0.93±0.4 | 0.92±0.22 |
| Pinocarvone | 14.10 | - | - | ND | ND | ND | ND |
| *endo*-Borneol | 14.18 | 2.29 | 0.164 | 0.48±0.16 | 1.2±0.72 | 2.85±1.26 | 2.2±0.53 |
| 3-Thujene-2-one | 14.34 | - | - | ND | ND | 0.02±0 | ND |
| Isopinocamphone | 14.40 | 0.18 | 0.681 | 1.3±0.68(a) | 0.91±0.54(a) | 1.27±0.62(a) | 1.36±0.02(a) |
| Terpinen-4-ol | 14.46 | 1.58 | 0.249 | 0.24±0.15(a) | 0.07±0.01(a) | 0.04±0(a) | 0.04±0.01(a) |
| ***p*-Cymene-8-ol** | 14.65 | 8.95 | **0.03** | 0.03±0(a) | 0.02±0(a) | 0.43±0.06(a) | 0.15±0.09(a) |
| α-Terpineol | 14.79 | 0.73 | 0.416 | 0.67±0.34 | 0.29±0.14 | 0.27±0.04 | 0.37±0.22 |
| Myrtenol | 14.94 | - | - | ND | ND | 0.4±0.2 | 0.32±0.2 |
| Verbenone | 15.28 | - | - | ND | ND | ND | ND |
| 2-Hydroxycineole | 15.58 | - | - | ND | ND | TR | TR |
| Thymol methyl ether | 15.85 | 0.66 | 0.436 | 1.03±0.21 | 0.67±0.23 | 0.54±0.22 | 0.76±0.17 |
| Myrtanol isomer1 | 16.73 | - | - | ND | ND | ND | ND |
| Myrtanol isomer2 | 16.30 | - | - | ND | ND | ND | ND |
| *p*-Menth-2-en-7-ol | 16.42 | - | - | ND | ND | ND | ND |
| Myrtanol isomer3 | 16.48 | - | - | ND | ND | ND | ND |
| Myrtenyl acetate isomer1 | 17.43 | - | - | ND | ND | 0.87±0 | ND |
| Myrtenyl acetate isomer2 | 18.13 | - | - | ND | ND | ND | ND |
| ***Sesquiterpenes*** |  |  |  |  |  |  |  |
| **α-Longipinene** | 18.63 | 12.26 | **0.007** | 0.2±0.03(a) | 0.09±0.02(ab) | 0.03±0(b) | 0.04±0.02(b) |
| **Longicyclene** | 19.10 | 10.45 | **0.012** | 0.26±0.04(a) | 0.07±0.01(a) | 0.03±0.01(a) | 0.06±0(a) |
| **Longifolene** | 19.88 | 6.48 | **0.031** | 2.07±0.32(a) | 1.07±0.22(a) | 0.42±0.11(a) | 0.74±0.46(a) |
| **(*E*)-β-Caryophyllene** | 20.17 | 20.58 | **0.001** | 8.59±0.44(a) | 4.01±0.47(ab) | 1.47±0.26(b) | 1.61±1.02(b) |
| (*E*)-β-Caryophyllene (fungus) | 20.56 | - | - | ND | ND | ND | ND |
| (*E*)-β-Farnesene | 20.84 | - | - | ND | ND | ND | ND |
| **Humulene** | 20.90 | 18.59 | **0.002** | 3.1±0.36(a) | 1.6±0.24(ab) | 0.67±0.15(b) | 0.57±0.32(b) |
| Caryophyllene oxide | 23.56 | - | - | ND | ND | ND | ND |

^#^- Estimated retention time from GC-MS

***^$^-***Significant differences between time points are denoted by small letters (ANOVA, followed by Tukey’s test, *P<0.05)*

***Table S4***. Relative amounts (mean ± SE, N=5) of volatiles detected at various time periods after inoculation of fresh spruce bark with *E. polonica* (4, 8, and 12 days). Volatiles were collected on polydimethylsiloxane tubes for 2 hours and were subjected to GC-MS analysis (see materials and methods section for details). ND=not detected, NA=not analyzed, TR= trace amounts (<500 TIC counts)

| ***Compounds*** | **RT^#^** | **F*^$^*** | **P*^$^*** | ***E. polonica* peak area (*10^4^ TIC counts)** | | |
| --- | --- | --- | --- | --- | --- | --- |
|  |  |  |  | **4d** | **8d** | **12d** |
| ***Aliphatics*** |  | | | | | |
| 2-Butanone | 1.85 | - | - | NA | NA | NA |
| 2-Methyl-3-buten-2-ol | 1.93 | - | - | NA | NA | NA |
| Ethyl acetate | 1.95 | - | - | NA | NA | NA |
| Isobutanol | 2.40 | 0.89 | 0.388 | TR | 0.14±0.03 | 0.26±0.08 |
| **Isopropyl acetate** | 2.33 | 27.91 | **<0.001** | 0.05±0.01(b) | 1.17±0.33(b) | 3.43±0.32(a) |
| **Acetoin** | 2.85 | 9.89 | **0.009** | 1.48±0.89(b) | 3.59±1.71(ab) | 10.7±4.04(a) |
| **Ethyl propanoate** | 2.88 | 16.04 | **<0.001** | 3.08±0.72(b) | 18.11±2.91(a) | 12.51±6.25(a) |
| **3-Methyl-1-butanol** | 3.24 | 18.87 | **<0.001** | 1.29±0.32(b) | 5.16±0.96(a) | 5.83±0.91(a) |
| **Ethyl isobutyrate** | 3.69 | 15.63 | **<0.001** | 0.38±0.18(b) | 1.45±0.46(ab) | 3.22±0.04(a) |
| **Isobutyl acetate** | 3.99 | 5.18 | **0.044** | 0.2±0.05(a) | 2.64±0.52(a) | 3.12±1.18(a) |
| 2,3-Butanediol | 4.17 | - | - | ND | ND | 0.05±0.01 |
| Ethyl butanoate | 4.55 | 0.9 | 0.366 | 0.54±0.16 | 10.41±2.08 | 4.22±1.49 |
| Ethyl but-2-enoate | 5.60 | 1.08 | 0.329 | 0.1±0.03 | 1.1±0.24 | 0.6±0.2 |
| Ethyl 2-methylbutyrate | 5.75 | 2.44 | 0.179 | 0.09±0.02 | 0.24±0.09 | ND |
| 1-Hexanol | 6.25 | - | - | ND | ND | ND |
| **3-Methyl-1-butyl acetate** | 6.46 | 12.04 | **0.005** | 22.15±4.34(a) | 14.24±1.55(ab) | 5.73±1.55(b) |
| Isopentyl-2-methylbutanoate | 12.47 | 0.16 | 0.858 | 0.45±0.25 | 0.23±0.11 | 0.15±0.06 |
| Isoamyl valerate | 12.60 | 1.43 | 0.289 | 1.33±0.69 | 0.49±0.28 | 0.18±0.11 |
| ***Aromatics*** |  | | | | | |
| 2-Phenylethyl alcohol | 12.79 | - | - | ND | ND | 0.07±0.01 |
| 2-Phenylethyl acetate | 16.39 | 0.59 | 0.461 | 0.21±0.05 | 0.38±0.11 | 0.29±0.04 |
| Citronellyl acetate | 18.58 | - | - | 0.85±0.18 | ND | 0.06±0.02 |
| ***Spiroketals*** |  | | | | | |
| *endo-*1,3-dimethyl-2,9-dioxabicyclo[3.3.1]nonane | 10.81 | - | - | ND | ND | ND |
| *trans*-Conophthorin | 11.29 | - | - | ND | ND | ND |
| Brevicomin | 11.64 | - | - | ND | ND | ND |
| *exo-*1,3-dimethyl-2,9-dioxabicyclo[3.3.1]nonane | 12.37 | - | - | ND | ND | TR |
| ***Monoterpenes*** |  | | | | | |
| **Santene** | 6.61 | 5.28 | **0.042** | 1.09±0.24(a) | 0.78±0.22(a) | 0.37±0.07(a) |
| **Tricyclene** | 7.67 | 11.03 | **0.007** | 3.55±0.49(a) | 2.46±0.91(ab) | 0.53±0.08(b) |
| **α-Thujene** | 7.76 | 16.15 | **<0.001** | 2.37±0.65(a) | 1.54±0.91(ab) | 0.22±0.11(b) |
| **α-Pinene** | 7.94 | 11.41 | **0.006** | 665±92.58(a) | 444±161.5(ab) | 114±15.67(b) |
| **Camphene** | 8.34 | 24.68 | **<0.001** | 10.86±1.25(a) | 8.37±3.08(a) | 2.04±0.28(b) |
| **Verbenene** | 8.51 | 13.88 | **<0.001** | 0.43±0.06(a) | 0.28±0.09(ab) | 0.07±0.01(b) |
| Sabinene | 9.50 | 1.66 | 0.245 | 0.81±0.41 | 0.37±0.21 | ND |
| **β-Pinene** | 9.13 | 16.24 | **<0.001** | 1186±133(a) | 728±274(ab) | 131±24.02(b) |
| **β-Myrcene** | 9.54 | 11.86 | **0.005** | 32.48±10.99(a) | 21.12±12.68(ab) | 3.02±1.47(b) |
| α-Phellandrene | 9.88 | 1.05 | 0.353 | 1.56±0.45 | 0.79±0.25 | ND |
| α-Terpinene | 10.21 | 1.6 | 0.253 | 0.32±0.13 | 0.35±0.15 | ND |
| ***p*-Cymene** | 10.43 | 9.23 | **0.011** | 18.83±3.69(a) | 11.63±3.88(ab) | 3.93±0.7(b) |
| **Limonene** | 10.51 | 22.34 | **<0.001** | 55.65±10.3(a) | 36.45±16.17(a) | 7.61±1.44(b) |
| **β-Phellandrene** | 10.55 | 13.93 | **<0.001** | 151±36.63(a) | 104±55.09(ab) | 19.93±7.27(b) |
| γ-Terpinene | 11.37 | - | - | 0.92±0.31 | ND | ND |
| α-Terpinolene | 12.16 | 1.29 | 0.308 | 1.07±0.27 | 0.82±0.33 | ND |
| *p*-Cymenene | 12.19 | 0.76 | 0.402 | 0.33±0.08 | 0.26±0.09 | 0.23±0.03 |
| ***Oxygenated monoterpenes*** |  | | | | | |
| **1,8-Cineole** | 10.61 | 8.66 | **0.013** | 4.45±0.7(a) | 3.19±0.61(ab) | 1.69±0.44(b) |
| Linalool oxide | 11.73 | - | - | TR | TR | ND |
| Fenchone | 12.15 | 1.68 | 0.222 | 0.52±0.11 | 0.48±0.18 | 0.25±0.1 |
| *trans*-4-Thujanol | 12.42 | - | - | ND | ND | ND |
| ***exo*-Fenchol** | 12.82 | 9.49 | **0.012** | 0.15±0.07(a) | 0.22±0.02(a) | 0.4±0.04(a) |
| Thujone | 12.93 | - | - | 0.04±0 | ND | ND |
| *p*-Isopropylcyclohexanol | 13.41 | - | - | ND | ND | 0.06±0.01 |
| *trans*-Pinocarveol | 13.48 | 3.02 | 0.11 | 0.37±0.16 | 0.49±0.09 | 0.63±0.11 |
| Camphor | 13.63 | 0.11 | 0.751 | 0.21±0.08 | 0.16±0.03 | 0.13±0.03 |
| Camphene hydrate | 13.73 | 0.43 | 0.663 | 0.1±0.05 | 0.12±0.01 | 0.16±0.03 |
| Pinocamphone | 14.43 | 3.39 | 0.093 | 2.5±0.45 | 2.2±0.23 | 1.45±0.09 |
| Pinocarvone | 14.10 | - | - | ND | ND | ND |
| *endo*-Borneol | 14.18 | 1.03 | 0.333 | 3.21±0.6 | 3.26±0.58 | 4.22±0.69 |
| 3-Thujene-2-one | 14.34 | - | - | ND | ND | ND |
| Isopinocamphone | 14.40 | 0.09 | 0.766 | 1.11±0.33 | 0.9±0.05 | 1.03±0.19 |
| Terpinen-4-ol | 14.46 | 0.28 | 0.608 | 0.97±0.43 | 0.64±0.23 | 0.78±0.19 |
| ***p*-Cymene-8-ol** | 14.65 | 29.44 | **<0.001** | 0.15±0.05 | 0.41±0.05 | 1.07±0.26 |
| α-Terpineol | 14.79 | 0.34 | 0.574 | 1.49±0.56 | 1.45±0.11 | 1.86±0.26 |
| **Myrtenol** | 14.94 | 41.22 | **<0.001** | ND | 0.65±0.12(b) | 2.72±0.35(a) |
| Verbenone | 15.28 | - | - | ND | ND | ND |
| 2-Hydroxycineole | 15.58 | - | - | ND | TR | TR |
| **Thymol methyl ether** | 15.85 | 5.65 | **0.037** | 2.37±0.76(a) | 1.52±0.28(a) | 0.83±0.14(a) |
| Myrtanol isomer1 | 16.73 | - | - | ND | ND | ND |
| Myrtanol isomer2 | 16.30 | - | - | ND | ND | ND |
| *p*-Menth-2-en-7-ol | 16.42 | - | - | ND | ND | ND |
| Myrtanol isomer3 | 16.48 | - | - | ND | ND | ND |
| Myrtenyl acetate isomer1 | 17.43 | 0.02 | 0.907 | 0.93±0.34 | 0.83±0.32 | ND |
| Myrtenyl acetate isomer2 | 18.13 | 0.39 | 0.549 | 0.79±0.22 | 0.43±0.1 | 0.68±0.15 |
| ***Sesquiterpenes*** |  | | | | | |
| **α-Longipinene** | 18.63 | 8.67 | **0.013** | 0.57±0.2(a) | 0.19±0.05(ab) | 0.09±0.04(b) |
| **Longicyclene** | 19.10 | 6.52 | **0.027** | 0.64±0.23(a) | 0.23±0.06(a) | 0.11±0.05(a) |
| **Longifolene** | 19.88 | 10.03 | **0.009** | 4.45±1.3(a) | 1.74±0.31(ab) | 1.08±0.26(b) |
| **(*E*)-β-Caryophyllene** | 20.17 | 20.48 | **<0.001** | 12.86±3.85(a) | 4.01±0.21(b) | 2.28±0.47(b) |
| (*E*)-β-Caryophyllene (fungus) | 20.56 | - | - | ND | ND | ND |
| **(*E*)-β-Farnesene** | 20.84 | 4.6 | **0.099** | 0.78±0.15 | 0.16±0.02 | ND |
| **Humulene** | 20.90 | 12.25 | **0.005** | 4.45±1.26(a) | 1.64±0.15(ab) | 0.99±0.1(b) |
| Caryophyllene oxide | 23.56 | - | - | ND | ND | ND |

^#^- Estimated retention time from GC-MS

***^$^-***Significant differences between time points are denoted by small letters (ANOVA, followed by Tukey’s test, *P<0.05)*

***Table S5***. Relative amounts (mean ± SE, N=5) of volatiles detected at various time periods after inoculation of fresh spruce bark with *G. penicillata* (4, 8, 12 and 18 days). Volatiles were collected on polydimethylsiloxane tubes for 2 hours and were subjected to GC-MS analysis (see materials and methods section for details). ND=not detected, NA=not analyzed, TR= trace amounts (<500 TIC counts)

| ***Compounds*** | **RT^#^** | **F*^$^*** | **P*^$^*** | **G. penicillata peak area (*10^4^ TIC counts)** | | | |
| --- | --- | --- | --- | --- | --- | --- | --- |
|  |  |  |  | **4d** | **8d** | **12d** | **18d** |
| ***Aliphatics*** |  | | | | | | |
| 2-Butanone | 1.85 | 0.01 | 0.914 | 4.02±0.94 | 3.8±0.81 | 6.83±3.55 | 1±0.2 |
| **2-Methyl-3-buten-2-ol** | 1.93 | 12.6 | **0.004** | 0.1±0.04(b) | 0.2±0.04(b) | 0.57±0.14(ab) | 0.86±0.18(a) |
| Ethyl acetate | 1.95 | 0.67 | 0.425 | 4.33±0.73 | 9.41±2.76 | 9.39±2.35 | 6.16±3.02 |
| Isobutanol | 2.40 | 0.81 | 0.386 | 0.51±0.07 | 1.11±0.19 | 1.24±0.03 | 0.66±0.28 |
| Isopropyl acetate | 2.33 | - | - | ND | ND | ND | ND |
| Acetoin | 2.85 | 0.6 | 0.449 | 5.21±2.25 | 5.06±0.54 | 10.21±3.15 | 8.16±5.12 |
| Ethyl propanoate | 2.88 | 1.97 | 0.184 | 1.79±0.6 | 0.3±0.11 | 0.25±0.09 | 0.34±0.13 |
| 3-Methyl-1-butanol | 3.24 | 0.03 | 0.871 | 4.52±0.79 | 12.8±0.37 | 8.47±2.56 | 5.11±2.46 |
| Ethyl isobutyrate | 3.69 | 2.57 | 0.184 | 0.77±0.13 | 0.35±0.08 | ND | ND |
| Isobutyl acetate | 3.99 | 0.07 | 0.8 | 0.29±0.07 | 1.19±0.37 | 0.93±0.42 | 0.65±0.35 |
| 2,3-Butanediol | 4.17 | 1.11 | 0.323 | ND | 0.09±0.01 | 0.23±0.03 | 0.16±0.04 |
| Ethyl butanoate | 4.55 | - | - | 0.49±0.21 | ND | ND | ND |
| Ethyl but-2-enoate | 5.60 | 0.14 | 0.721 | 0.25±0.06 | 0.38±0.03 | 0.5±0.07 | 0.29±0 |
| Ethyl 2-methylbutyrate | 5.75 | - | - | 0.09±0.02 | ND | ND | ND |
| 1-Hexanol | 6.25 | 0.7 | 0.422 | 0.19±0.06 | 0.78±0.28 | 0.83±0.46 | ND |
| **3-Methyl-1-butyl acetate** | 6.46 | 6.19 | **0.027** | 0.43±0.12(a) | 1.55±0.49(a) | 1.28±0.19(a) | 1.34±0.14(a) |
| Isopentyl-2-methylbutanoate | 12.47 | 0.6 | 0.594 | 0.13±0.04 | 0.08±0.02 | 0.05±0.01 | ND |
| Isoamyl valerate | 12.60 | 1.28 | 0.338 | 0.37±0.11 | 0.17±0.07 | 0.11±0.03 | 0.08±0.03 |
| ***Aromatics*** |  | | | | | | |
| 2-Phenylethyl alcohol | 12.79 | 2.19 | 0.167 | ND | 0.73±0.11 | 0.94±0.17 | 1.19±0.25 |
| 2-Phenylethyl acetate | 16.39 | 0.33 | 0.577 | 0.19±0.05 | 0.34±0.11 | 0.07±0.02 | ND |
| Citronellyl acetate | 18.58 | - | - | ND | ND | ND | ND |
| ***Spiroketals*** |  | | | | | | |
| *endo-*1,3-dimethyl-2,9-dioxabicyclo[3.3.1]nonane | 10.81 | 0.4 | 0.537 | 0.2±0.07 | 0.12±0.04 | 0.07±0.01 | 0.11±0.02 |
| *trans*-Conophthorin | 11.29 | 1.9 | 0.191 | 0.03±0 | 0.9±0.46 | 0.84±0.37 | 0.75±0.32 |
| Brevicomin | 11.64 |  |  | ND | ND | ND | ND |
| ***exo-*1,3-dimethyl-2,9-dioxabicyclo[3.3.1]nonane** | 12.37 | 7.1 | **0.017** | 1.91±0.48(a) | 1.02±0.28(a) | 0.54±0.1(a) | 0.52±0.22(a) |
| ***Monoterpenes*** |  | | | | | | |
| Santene | 6.61 | 1.83 | 0.194 | 0.96±0.25 | 0.54±0.18 | 0.42±0.09 | 0.49±0.11 |
| **Tricyclene** | 7.67 | 25.75 | **<0.001** | 3.31±0.88(a) | 0.9±0.31(ab) | 0.4±0.11(b) | 0.27±0.09(b) |
| **α-Thujene** | 7.76 | 26.45 | **<0.001** | 5.41±2.65(a) | 1.33±0.6(ab) | 0.43±0.16(b) | 0.18±0.06(b) |
| **α-Pinene** | 7.94 | 42.14 | **<0.001** | 683±205(a) | 193±69(ab) | 82.4±23.45(ab) | 36.17±10(c) |
| **Camphene** | 8.34 | 20.99 | **<0.001** | 11.08±2.91(a) | 3.01±0.89(ab) | 1.35±0.45(b) | 1.16±0.44(b) |
| **Verbenene** | 8.51 | 14.53 | **0.001** | 0.79±0.45(a) | 0.14±0.05(ab) | 0.06±0.01(b) | 0.05±0.01(b) |
| Sabinene | 9.50 |  |  | 0.32±0.03 | ND | ND | ND |
| **β-Pinene** | 9.13 | 136.93 | **<0.001** | 800±108(a) | 159±19.35(b) | 45.92±9.33(c) | 13.28±4.35(d) |
| **β-Myrcene** | 9.54 | 140.32 | **<0.001** | 22.45±4.99(a) | 6.26±1.06(b) | 1.98±0.23(c) | 0.77±0.07© |
| **Unknown** | 9.85 | 7.3 | **0.003** | 3.57±0.95(a) | 0.27±0.07(b) | 0.28±0.18(b) | 0.91±0.34(ab) |
| **α-Phellandrene** | 9.88 | 13.53 | **0.002** | 1.82±0.53(a) | 0.8±0.35(ab) | 0.28±0.13(b) | 0.27±0.14(b) |
| **α-Terpinene** | 10.21 | 9.77 | **0.007** | 0.6±0.26(a) | 0.25±0.11(ab) | 0.09±0.02(ab) | 0.07±0.02(b) |
| *p*-Cymene | 10.43 | 2.2 | 0.156 | 42.27±20.53 | 24.38±11.35 | 13.31±5.88 | 16.3±6.77 |
| **Limonene** | 10.51 | 10.33 | **0.005** | 56.38±19.04(a) | 19.68±7.25(ab) | 10.11±3.55(b) | 10.27±3.71(b) |
| **β-Phellandrene** | 10.55 | 25.35 | **<0.001** | 180±51.67(a) | 70.17±28.06(ab) | 25.49±9.81(b) | 17.26±6.77(b) |
| **1,8-Cineole** | 10.61 | 40.56 | **<0.001** | 7.18±1.06(a) | 3.57±0.45(ab) | 1.38±0.3(bc) | 0.79±0.23(c) |
| **γ-Terpinene** | 11.37 | 11.51 | **0.004** | 2.43±1.15(a) | 1.32±0.4(ab) | 0.3±0.08(b) | 0.33±0.08(ab) |
| ***Oxygenated monoterpenes*** |  | | | | | | |
| **Linalool oxide** | 11.73 | 13.25 | **0.002** | 0.19±0.06(b) | 0.4±0.08(ab) | 1.06±0.34(a) | 1.38±0.38(a) |
| **Fenchone** | 12.15 | 10.96 | **0.004** | 2.2±1.04(b) | 2.63±1.33(b) | 10.74±5.68(b) | 24.42±4.58(a) |
| α-Terpinolene | 12.16 | 1.34 | 0.263 | 6.06±4.09 | 2.8±1.43 | 1.02±0.29 | 1.61±0.52 |
| *p*-Cymenene | 12.19 | 0.72 | 0.407 | 2.07±1.22 | 2.14±0.85 | 2.92±1.77 | 5.07±3 |
| *trans*-4-Thujanol | 12.42 | - | - | ND | ND | ND | TR |
| *exo*-Fenchol | 12.82 | 2.7 | 0.119 | 0.67±0.45 | 1.44±0.55 | 1.78±0.36 | 1.42±0.63 |
| Thujone | 12.93 | 2.8 | 0.123 | 0.11±0.03 | 0.07±0.02 | 0.05±0 | 0.05±0.01 |
| *p*-Isopropylcyclohexanol | 13.41 | 4.1 | 0.064 | 0.32±0.12 | 0.23±0.09 | 0.43±0.17 | 1.11±0.4 |
| ***trans*-Pinocarveol** | 13.48 | 7.18 | **0.016** | 0.59±0.2(a) | 1.44±0.29(a) | 2.68±0.48(a) | 2.36±0.66(a) |
| **Camphor** | 13.63 | 13.06 | **0.002** | 21.83±6.85(b) | 21.02±7.15(b) | 74.89±32.59(ab) | 148±6.25(a) |
| Camphene hydrate | 13.73 | 0.78 | 0.524 | 0.25±0.14 | 0.38±0.14 | 0.27±0.08 | 0.33±0.07 |
| Pinocamphone | 14.43 | 0.13 | 0.727 | 4.52±0.61 | 3.51±0.66 | 3.15±0.82 | 4.07±0.91 |
| Pinocarvone | 14.10 | - | - | ND | ND | ND | ND |
| endo-Borneol | 14.18 | 1.29 | 0.275 | 4.34±0.61 | 9.52±1.35 | 11.67±4.18 | 6.46±2.68 |
| 3-Thujene-2-one | 14.34 | - | - | ND | ND | ND | ND |
| Isopinocamphone | 14.40 | 3.72 | 0.071 | 3.12±1.07(b) | 2.97±1.01(ab) | 5.43±2.12(a) | 10.59±2.84(a) |
| Terpinen-4-ol | 14.46 | 0.59 | 0.455 | 23.53±16.33(a) | 56.28±22.27(a) | 56.52±19.3(a) | 47.27±14.19(a) |
| ***p*-Cymene-8-ol** | 14.65 | 12.45 | **0.003** | 1.02±0.51(b) | 2.52±0.63(ab) | 2.99±0.48(ab) | 3.84±0.52(a) |
| α-Terpineol | 14.79 | 3.96 | 0.063 | 7.6±2.93 | 12.84±4.13 | 20.09±5.84 | 20.15±4.63 |
| **Myrtenol** | 14.94 | 16.37 | **0.001** | 4.42±1.22(b) | 15.32±2.87(a) | 24.84±5.61(a) | 35.57±10.83(a) |
| Verbenone | 15.28 | 0.18 | 0.679 | ND | 0.04±0.01 | 0.05±0 | 0.03±0.01 |
| **2-Hydroxycineole** | 15.58 | 10.48 | **0.008** | 0.05±0.01(b) | 0.11±0.01(ab) | 0.18±0.05(ab) | 0.25±0.03(a) |
| **Thymol methyl ether** | 15.85 | 7.33 | **0.015** | 2.88±0.76(a) | 2.04±0.67(a) | 1.25±0.46(a) | 1.09±0.43(a) |
| **Myrtanol isomer1** | 16.73 | 14.04 | **0.005** | ND | 0.05±0.01(b) | 0.16±0.06(ab) | 0.28±0.06(a) |
| **Myrtanol isomer2** | 16.30 | 18.09 | **0.001** | 0.04±0.01(c) | 0.1±0.03(bc) | 0.38±0.14(ab) | 0.79±0.18(a) |
| *p*-Menth-2-en-7-ol | 16.42 | - | - | ND | TR | TR | TR |
| **Myrtanol isomer3** | 16.48 | 10.12 | **0.011** | ND | 0.19±0.05(b) | 0.61±0.13(ab) | 1.03±0.36(a) |
| **Myrtenyl acetate isomer1** | 17.43 | 8.37 | **0.02** | 1.9±0.46(a) | 0.13±0.03(a) | 0.27±0.14(a) | ND |
| Myrtenyl acetate isomer2 | 18.13 | - | - | ND | ND | 0.05±0 | ND |
| ***Sesquiterpenes*** |  | | | | | | |
| α-Longipinene | 18.63 | 3.23 | 0.097 | 0.39±0.16 | 0.15±0.08 | 0.1±0.04 | 0.11±0.04 |
| **Longicyclene** | 19.10 | 9.4 | **0.007** | 0.44±0.16(a) | 0.23±0.11(a) | 0.11±0.06(a) | 0.09±0.05(a) |
| **Longifolene** | 19.88 | 9.65 | **0.006** | 2.96±0.86(a) | 1.69±0.73(ab) | 0.85±0.48(ab) | 0.69±0.41(b) |
| **(*E*)-β-Caryophyllene** | 20.17 | 18.57 | **<0.001** | 9.7±2.05(a) | 5.52±1.62(ab) | 2.2±0.94(b) | 1.87±0.84(b) |
| (*E*)-β-Caryophyllene (fungus) | 20.56 | 0 | 0.992 | ND | 0.51±0.15 | 0.44±0.15 | 0.8±0.43 |
| **(*E*)-β-Farnesene** | 20.84 | 12.11 | **0.01** | 0.36±0.05(a) | 0.12±0.02(b) | ND | ND |
| **Humulene** | 20.90 | 10.57 | **0.005** | 3±0.5(a) | 1.94±0.59(ab) | 1.05±0.41(ab) | 0.97±0.39(b) |
| Caryophyllene oxide | 23.56 | - | - | ND | 0.7±0.12 | 0.66±0.12 | 0.86±0.18 |

^#^- Estimated retention time from GC-MS

***^$^-***Significant differences between time points are denoted by small letters (ANOVA, followed by Tukey’s test, *P<0.05)*

***Table S6.*** Relative amounts (mean ± SE, N=5) of volatiles detected at various time periods after inoculation of fresh spruce bark with *L. europhioides* (4, 8, 12 and 18 days). Volatiles were collected on polydimethylsiloxane tubes for 2 hours and were subjected to GC-MS analysis (see materials and methods section for details). ND=not detected, NA=not analyzed, TR= trace amounts (<500 TIC counts)

| ***Compounds*** | **RT^#^** | **F*^$^*** | **P*^$^*** | ***L. europhioides* peak area (*10^4^ TIC counts)** | | | |
| --- | --- | --- | --- | --- | --- | --- | --- |
|  |  |  |  | **4d** | **8d** | **12d** | **18d** |
| ***Aliphatics*** |  | | | | | | |
| **2-Butanone** | 1.85 | 5.86 | **0.027** | 3.06±0.69(a) | NA | 2.27±1.13(a) | 3.04±0.65(a) |
| **2-Methyl-3-buten-2-ol** | 1.93 | 42.57 | **<0.001** | 0.59±0.09(b) | 4.35±2.05(a) | 11.13±3.29(a) | 13.82±2.62(a) |
| **Ethyl acetate** | 1.95 | 14.65 | **0.002** | 3.38±0.79(a) | 0.57±0.34(ab) | 0.21±0.11(b) | 0.23±0.14(b) |
| Isobutanol | 2.40 | 1.64 | 0.227 | 2.5±0.23(ab) | 5.94±1.52(a) | 1.13±0.39(b) | TR |
| Isopropyl acetate | 2.33 | - | - | ND | ND | ND | ND |
| Acetoin | 2.85 | 4.48 | 0.051 | 12.69±3.21 | 8.92±2.51 | 7.86±3.25 | 2.39±1.2 |
| Ethyl propanoate | 2.88 | 0.03 | 0.877 | 0.27±0.16 | TR | ND | 0.13±0.03 |
| **3-Methyl-1-butanol** | 3.24 | 13.94 | **0.002** | 14.04±1.12(a) | 20.8±1.32(ab) | 8.66±3.5(bc) | 1.02±0.3 (c) |
| Ethyl isobutyrate | 3.69 | - | - | 0.62±0.3 | ND | ND | ND |
| Isobutyl acetate | 3.99 | 2.11 | 0.18 | 0.54±0.14 | 0.38±0.04 | 0.26±0.06 | ND |
| 2,3-Butanediol | 4.17 | 0.44 | 0.52 | 0.11±0 | 0.09±0.03 | 0.16±0.06 | 0.15±0.05 |
| Ethyl butanoate | 4.55 | - | - | ND | ND | ND | ND |
| Ethyl but-2-enoate | 5.60 | 0.05 | 0.831 | 0.78±0.09 | 1.4±0.21 | 0.54±0.23 | ND |
| Ethyl 2-methylbutyrate | 5.75 | - | - | ND | ND | ND | ND |
| **1-Hexanol** | 6.25 | 8.81 | **0.021** | 1.07±0.17(b) | 1.95±0.18(a) | ND | ND |
| 3-Methyl-1-butyl acetate | 6.46 | 4.26 | 0.061 | 0.16±0.03(b) | 0.24±0.04(b) | 0.2±0.1(ab) | 1.19±0.11(a) |
| Isopentyl-2-methylbutanoate | 12.47 | - | - | 1.08±0.46 | 0.71±0.3 | ND | ND |
| Isoamyl valerate | 12.60 | 0.13 | 0.88 | 1.02±0.8 | 1.12±0.58 | ND | ND |
| ***Aromatics*** |  | | | | | | |
| 2-Phenylethyl alcohol | 12.79 | 1.13 | 0.303 | 0.61±0.11 | 1.64±0.3 | 1.2±0.14 | 1.22±0.18 |
| 2-Phenylethyl acetate | 16.39 | - | - | ND | ND | ND | ND |
| Citronellyl acetate | 18.58 | - | - | ND | ND | ND | ND |
| ***Spiroketals*** |  | | | | | | |
| *endo-*1,3-dimethyl-2,9-dioxabicyclo[3.3.1]nonane | 10.81 | 4.06 | 0.061 | 1.31±0.24 | 1.02±0.2 | 0.78±0.12 | 0.75±0.12 |
| ***trans*-Conophthorin** | 11.29 | 69.34 | **<0.001** | 0.03±0(c) | 0.05±0(b) | 0.06±0.01(b) | 0.15±0.02(a) |
| Brevicomin | 11.64 | - | - | TR | TR | ND | TR |
| ***exo-*1,3-dimethyl-2,9-dioxabicyclo[3.3.1]nonane** | 12.37 | 3.77 | **0.07** | 2.28±0.37(a) | 1.77±0.29(a) | 1.19±0.04(a) | 1.52±0.12(a) |
| ***Monoterpenes*** |  | | | | | | |
| Santene | 6.61 | 1.83 | 0.194 | 0.97±0.27 | 0.93±0.17 | 0.48±0.16 | 0.46±0.05 |
| **Tricyclene** | 7.67 | 48.16 | **<0.001** | 3.15±0.46(a) | 0.89±0.26(b) | 0.31±0.12(bc) | 0.14±0.03(c) |
| **α-Thujene** | 7.76 | 21.98 | **<0.001** | 3.17±1.01(a) | 0.73±0.33(ab) | 0.18±0.11(b) | 0.11±0.06(b) |
| **α-Pinene** | 7.94 | 56.3 | **<0.001** | 633±111(a) | 189±46.47(b) | 60.23±18.76(bc) | 33±7.3(c) |
| **Camphene** | 8.34 | 49.67 | **<0.001** | 10.4±1.71(a) | 3.7±1.03(b) | 1.33±0.41(bc) | 0.72±0.19(c) |
| **Verbenene** | 8.51 | 22.45 | **<0.001** | 0.44±0.05(a) | 0.15±0.05(ab) | 0.06±0.02(b) | 0.05±0.01(b) |
| Sabinene | 9.50 | - | - | 0.22±0.07 | ND | ND | ND |
| **β-Pinene** | 9.13 | 89.32 | **<0.001** | 907±161(a) | 208±58.89(b) | 45.89±13.38(c) | 14.74±3.6(c) |
| **β-Myrcene** | 9.54 | 22.04 | **<0.001** | 28.05±11.54(a) | 10.09±4.59(ab) | 1.68±0.96(bc) | 1.28±0.9(c) |
| **Unknown** | 9.85 | 4.56 | **0.02** | 3.92±1.26(a) | 1.88±0.63(ab) | 0.66±0.26(ab) | 0.36±0.14(b) |
| **α-Phellandrene** | 9.88 | 4.94 | **0.05** | 1.34±0.4(a) | 0.87±0.27(a) | 0.12±0.04(a) | ND |
| α-Terpinene | 10.21 | 3 | 0.118 | 0.32±0.1 | 0.28±0.13 | 0.05±0.01 | ND |
| ***p*-Cymene** | 10.43 | 18.43 | **<0.001** | 30.27±6.76(a) | 11.94±2.98(ab) | 4.54±1.74(b) | 4.15±1.55(b) |
| **Limonene** | 10.51 | 27.95 | **<0.001** | 39.56±12.8(a) | 17.23±6.19(ab) | 4.67±2.18(bc) | 2.66±1.37(c) |
| **β-Phellandrene** | 10.55 | 22.57 | **<0.001** | 169±63.07(a) | 70.29±30.68(ab) | 13.38±6.65(bc) | 10.7±7.03(c) |
| γ-Terpinene | 11.37 | 3.2 | 0.099 | 1.25±0.34(a) | 0.99±0.45(ab) | 0.28±0.17(b) | 0.39±0.17(ab) |
| α-Terpinolene | 12.16 | 2.39 | 0.141 | 2.84±1.09 | 3.41±1.7 | 1.17±0.73 | 1.37±0.9 |
| *p*-Cymenene | 12.19 | 0.56 | 0.464 | 1.44±0.44 | 1.38±0.28 | 1.25±0.5 | 1.92±0.59 |
| ***Oxygenated monoterpenes*** |  | | | | | | |
| **1,8-Cineole** | 10.61 | 15.84 | **0.001** | 3.43±0.9(a) | 1.92±0.75(ab) | 0.35±0.04(b) | 0.72±0.4(b) |
| **Linalool oxide** | 11.73 | 4.17 | **0.058** | 0.3±0.08(a) | 0.81±0.16(a) | 0.6±0.08(a) | 0.97±0.31(a) |
| **Fenchone** | 12.15 | 13.12 | **0.002** | 1.97±0.47(b) | 2.15±0.43(b) | 4.99±1.82(b) | 16.14±2.56(a) |
| *trans*-4-Thujanol | 12.42 | - | - | ND | ND | ND | TR |
| ***exo*-Fenchol** | 12.82 | 4.99 | **0.04** | 1.47±0.27(b) | 3.13±0.56(ab) | 3.03±0.79(ab) | 3.21±0.32(a) |
| Thujone | 12.93 | 1.77 | 0.21 | 0.06±0.02 | 0.05±0.02 | 0.03±0 | 0.03±0 |
| *p*-Isopropylcyclohexanol | 13.41 | 0.08 | 0.785 | 0.48±0.12 | 0.9±0.13 | 0.94±0.27 | 1.04±0.68 |
| ***trans*-Pinocarveol** | 13.48 | 14.4 | **0.003** | 1.08±0.08(a) | 1.27±0.18(a) | 0.05±0.01(b) | ND |
| **Camphor** | 13.63 | 69.66 | **<0.001** | 10.13±1.79(c) | 57.6±6.22(b) | 73.08±17.48(ab) | 110±11.07(a) |
| Camphene hydrate | 13.73 | 1.26 | 0.324 | 0.41±0.09 | 0.74±0.14 | 0.42±0.15 | 0.61±0.11 |
| Pinocamphone | 14.43 | 0.83 | 0.375 | 4.84±0.4 | 4.44±0.41 | 3.58±0.87 | 6.1±0.56 |
| Pinocarvone | 14.10 | - | - | ND | ND | ND | ND |
| ***endo*-Borneol** | 14.18 | 94.37 | **<0.001** | 22.06±1.38(a) | 18.23±4.28(a) | 1.46±0.27(b) | 0.25±0.03(c) |
| 3-Thujene-2-one | 14.34 | 2.37 | 0.143 | 0.36±0.06 | 0.78±0.15 | 0.63±0.05 | 0.7±0.06 |
| **Isopinocamphone** | 14.40 | 29.4 | **<0.001** | 5.84±0.87(b) | 16.38±1.86(ab) | 22.14±2.88(a) | 25.91±3.26(a) |
| Terpinen-4-ol | 14.46 | 0.24 | 0.632 | 1.37±0.63 | 7.5±3.74 | 0.96±0.33 | ND |
| ***p*-Cymene-8-ol** | 14.65 | 5.46 | **0.033** | 1.34±0.3(a) | 1.87±0.32(a) | 1.72±0.5(a) | 2.71±0.57(a) |
| α-Terpineol | 14.79 | 0.35 | 0.563 | 10.21±1.21 | 16.17±3.82 | 13.92±4.02 | 7.96±1.35 |
| Myrtenol | 14.94 | 1.03 | 0.325 | 11.75±1.51 | 23.25±3.76 | 19.22±5.36 | 18.48±3.96 |
| Verbenone | 15.28 | 2.05 | 0.176 | 0.15±0.04 | 0.14±0.06 | 0.15±0.03 | 0.33±0.14 |
| **2-Hydroxycineole** | 15.58 | 16.43 | **0.001** | 0.1±0.01(b) | 0.35±0.05(a) | 0.3±0.04(ab) | 0.48±0.03(a) |
| **Thymol methyl ether** | 15.85 | 6.37 | **0.022** | 2.79±0.94(a) | 2.25±0.94(a) | 0.57±0.18(a) | 1.2±0.72(a) |
| **Myrtanol isomer1** | 16.73 | 12.26 | **0.005** | ND | 0.1±0.02(b) | 0.25±0.05(ab) | 0.38±0.08(a) |
| **Myrtanol isomer2** | 16.30 | 59.93 | **<0.001** | 0.09±0.02(b) | 0.28±0.06(b) | 0.68±0.06(a) | 1.02±0.16(a) |
| *p*-Menth-2-en-7-ol | 16.42 | - | - | TR | TR | TR | 0.05±0 |
| **Myrtanol isomer3** | 16.48 | 51.63 | **<0.001** | 0.13±0.03(c) | 0.48±0.11(bc) | 1.06±0.1(ab) | 1.62±0.25(a) |
| Myrtenyl acetate isomer1 | 17.43 | - | - | 0.52±0.21 | ND | ND | ND |
| Myrtenyl acetate isomer2 | 18.13 | - | - | ND | ND | ND | ND |
| ***Sesquiterpenes*** |  | | | | | | |
| **α-Longipinene** | 18.63 | 8.58 | **0.012** | 0.52±0.24(a) | 0.32±0.15(a) | 0.05±0.01(a) | 0.07±0.03(a) |
| **Longicyclene** | 19.10 | 4.61 | **0.049** | 0.62±0.28(a) | 0.37±0.19(a) | 0.07±0.02(a) | 0.16±0.08(a) |
| **Longifolene** | 19.88 | 6.88 | **0.018** | 3.54±1.3(a) | 2.91±1.3(a) | 0.58±0.23(a) | 1.09±0.65(a) |
| **(*E*)-β-Caryophyllene** | 20.17 | 7.18 | **0.016** | 11.02±3.28(a) | 8.75±3.2(a) | 2.15±0.73(a) | 3.1±1.52(a) |
| (*E*)-β-Caryophyllene (fungus) | 20.56 | 2.36 | 0.144 | 3.09±0.51 | 6.9±1.21 | 9.33±1.7 | 6.37±1.45 |
| (*E*)-β-Farnesene | 20.84 | 1.76 | 0.212 | 0.34±0.07 | 0.43±0.1 | 0.16±0.02 | 0.18±0.03 |
| Humulene | 20.90 | 3.98 | 0.062 | 3.77±0.94 | 3.2±1.11 | 1.06±0.35 | 1.67±0.79 |
| Caryophyllene oxide | 23.56 | - | - | ND | 0.25±0.05 | ND | ND |

^#^- Estimated retention time from GC-MS

***^$^-***Significant differences between time points are denoted by small letters (ANOVA, followed by Tukey’s test, *P<0.05)*

***Table S7***. Relative amounts (mean ± SE, N=5) of volatiles detected at various time periods after inoculation of fresh spruce bark with *O. bicolor* (4, 8, 12 and 18 days). Volatiles were collected on polydimethylsiloxane tubes for 2 hours and were subjected to GC-MS analysis (see materials and methods section for details). ND=not detected, NA=not analyzed, TR= trace amounts (<500 TIC counts)

| ***Compounds*** | **RT^#^** | **F*^$^*** | **P*^$^*** | ***O. bicolor* peak area (*10^4^ TIC counts)** | | | |
| --- | --- | --- | --- | --- | --- | --- | --- |
|  |  |  |  | **4d** | **8d** | **12d** | **18d** |
| ***Aliphatics*** |  | | | | | | |
| 2-Butanone | 1.85 | 0.25 | 0.621 | 3.05±0.82 | 2.58±0.33 | 2.83±0.74 | 2.82±1.2 |
| 2-Methyl-3-buten-2-ol | 1.93 | - | - | ND | ND | ND | ND |
| **Ethyl acetate** | 1.95 | 1.2 | 0.291 | 0.81±0.29 | 1.43±0.15 | 1.76±0.48 | 1.68±0.49 |
| **Isobutanol** | 2.40 | 13.83 | **0.002** | 0.5±0.119(b) | 1.92±0.2(a) | 2.37±0.3(a) | 1.34±0.51(a) |
| Isopropyl acetate | 2.33 | - | - | ND | ND | ND | ND |
| **Acetoin** | 2.85 | 26.35 | **<0.001** | 6.06±1.7(b) | 6.18±1.55(b) | 12.23±1.9(ab) | 25.51±4.6(a) |
| Ethyl propanoate | 2.88 | 3.64 | 0.079 | 1.17±0.45 | 0.51±0.25 | 0.2±0.07 | 0.08±0.01 |
| **3-Methyl-1-butanol** | 3.24 | 26.82 | **<0.001** | 7.49±0.85(b) | 18.51±0.81(a) | 26.51±1.21(a) | 27.05±3.47(a) |
| Ethyl isobutyrate | 3.69 | 1.21 | 0.3 | 1.23±0.39 | 2.23±0.86 | 0.29±0.09 | ND |
| **Isobutyl acetate** | 3.99 | 8.12 | **0.019** | ND | 0.17±0.01(b) | 0.17±0.03(b) | 0.46±0.03(a) |
| 2,3-Butanediol | 4.17 | 2.51 | 0.137 | 0.05±0.01 | 0.29±0.1 | 0.16±0.03 | 0.32±0.09 |
| Ethyl butanoate | 4.55 | 3.46 | 0.09 | 0.28±0.08 | 0.26±0.09 | 0.1±0.02 | 0.08±0.02 |
| Ethyl but-2-enoate | 5.60 | 0.74 | 0.409 | 0.3±0.05 | 0.6±0.22 | 0.25±0.02 | 0.21±0.01 |
| Ethyl 2-methylbutyrate | 5.75 | 0.45 | 0.527 | 0.18±0.07 | 0.27±0.08 | ND | ND |
| 1-Hexanol | 6.25 | 0.33 | 0.575 | 0.25±0.06 | 0.56±0.05 | 0.32±0.15 | 0.11±0.04 |
| **3-Methyl-1-butyl acetate** | 6.46 | 12.07 | **0.003** | 0.06±0.01(b) | 0.19±0.03(ab) | 0.2±0.03(a) | 0.35±0.12(a) |
| Isopentyl-2-methylbutanoate | 12.47 | 0.7 | 0.583 | 0.13±0.04 | 0.05±0.01 | 0.08±0.02 | 0.1±0.04 |
| Isoamyl valerate | 12.60 | 0.86 | 0.5 | 0.41±0.14 | 0.31±0.12 | 0.19±0.07 | 0.23±0.1 |
| ***Aromatics*** |  | | | | | | |
| 2-Phenylethyl alcohol | 12.79 | 0.97 | 0.341 | 0.34±0.09 | 0.55±0.12 | 0.64±0.1 | 0.58±0.15 |
| 2-Phenylethyl acetate | 16.39 | - | - | ND | ND | ND | ND |
| Citronellyl acetate | 18.58 | - | - | ND | ND | ND | ND |
| ***Spiroketals*** |  | | | | | | |
| *endo-*1,3-dimethyl-2,9-dioxabicyclo[3.3.1]nonane | 10.81 | 1.65 | 0.217 | 0.3±0.08 | 0.47±0.09 | 0.48±0.08 | 0.46±0.06 |
| *trans*-Conophthorin | 11.29 | - | - | ND | ND | ND | ND |
| Brevicomin | 11.64 | - | - | ND | ND | TR | TR |
| *exo-*1,3-dimethyl-2,9-dioxabicyclo[3.3.1]nonane | 12.37 | 3.63 | 0.074 | 0.41±0.07 | 1.01±0.17 | 0.85±0.12 | 0.91±0.13 |
| ***Monoterpenes*** |  | | | | | | |
| **Santene** | 6.61 | 19.19 | **<0.001** | 1.08±0.29(a) | 0.72±0.08(ab) | 0.41±0.06(b) | 0.36±0.02(b) |
| **Tricyclene** | 7.67 | 48.9 | **<0.001** | 4.35±0.82(a) | 1.69±0.26(a) | 0.59±0.19(b) | 0.35±0.07(b) |
| **α-Thujene** | 7.76 | 38.7 | **<0.001** | 2.15±0.61(a) | 0.73±0.16(ab) | 0.25±0.12(bc) | 0.12±0.05(c) |
| **α-Pinene** | 7.94 | 71.46 | **<0.001** | 744±118(a) | 293±35.35(b) | 120±25.59(c) | 75.79±11.16(c) |
| **Camphene** | 8.34 | 60.9 | **<0.001** | 14.64±2.78(a) | 5.56±1.03(a) | 2.07±0.62(b) | 1.07±0.22(b) |
| Verbenene | 8.51 | 3.44 | 0.088 | 0.77±0.29 | 0.62±0.24 | 0.37±0.18 | 0.19±0.07 |
| Sabinene | 9.50 | 0.67 | 0.46 | 0.73±0.3 | 0.13±0.07 | ND | ND |
| **β-Pinene** | 9.13 | 94.71 | **<0.001** | 1162±173(a) | 423±62.9(b) | 135±32.81(c) | 68.84±15.77(c) |
| **β-Myrcene** | 9.54 | 37.17 | **<0.001** | 24.6±6.54(a) | 11.2±1.83(ab) | 3.75±1.29(bc) | 1.88±0.89(c) |
| Unknown | 9.85 | 0.53 | 0.664 | 1.93±0.61 | 2.65±0.62 | 2.07±0.5 | 1.72±0.46 |
| **α-Phellandrene** | 9.88 | 16.52 | **0.002** | 1.6±0.4(a) | 0.74±0.19(ab) | 0.43±0.18(ab) | 0.2±0.05(b) |
| **α-Terpinene** | 10.21 | 12.61 | **0.005** | 0.4±0.169(a) | 0.16±0.04(ab) | 0.08±0.02(ab) | 0.03±0(b) |
| ***p*-Cymene** | 10.43 | 39.19 | **<0.001** | 26.89±5.3(a) | 11.6±2.53(ab) | 4.87±1.45(bc) | 3.04±0.52(c) |
| **Limonene** | 10.51 | 44.23 | **<0.001** | 51.15±10.26(a) | 21.07±3.56(a) | 7.58±2.15(b) | 4.79±0.94(b) |
| **β-Phellandrene** | 10.55 | 46.17 | **<0.001** | 152.±39.71(a) | 65.86±8.5(a) | 22.77±8.59(b) | 11.91±4.2(b) |
| γ-Terpinene | 11.37 | 0.86 | 0.422 | 1.36±0.36 | 0.67±0.01 | ND | ND |
| **α-Terpinolene** | 12.16 | 33.82 | **<0.001** | 1.05±0.23(a) | 0.73±0.12(a) | 0.21±0.06(b) | 0.16±0.04(b) |
| *p*-Cymenene | 12.19 | 0.37 | 0.551 | 0.59±0.15 | 0.57±0.11 | 0.57±0.13 | 0.88±0.36 |
| ***Oxygenated monoterpenes*** |  | | | | | | |
| **1,8-Cineole** | 10.61 | 47.05 | **<0.001** | 6.93±1.31(a) | 4.3±0.64(a) | 1.7±0.33(b) | 0.79±0.14(b) |
| Linalool oxide | 11.73 | 1.88 | 0.188 | 0.13±0.03 | 0.31±0.08 | 0.4±0.15 | 0.39±0.15 |
| **Fenchone** | 12.15 | 10.96 | **0.004** | 1.64±0.35(b) | 1.76±0.59(b) | 1.36±0.37(ab) | 2.26±1.07(a) |
| *trans*-4-Thujanol | 12.42 | - | - | ND | ND | ND | ND |
| ***exo*-Fenchol** | 12.82 | 33.56 | **<0.001** | 0.27±0.06(c) | 0.51±0.1(bc) | 0.68±0.08(ab) | 1.63±0.38(a) |
| **Thujone** | 12.93 | 6.79 | **0.022** | 0.24±0.08(a) | 0.12±0.05(a) | 0.06±0.02(a) | 0.04±0(a) |
| *p*-Isopropylcyclohexanol | 13.41 | 2.47 | 0.142 | ND | 0.15±0.03 | 0.19±0.06 | 0.33±0.1 |
| ***trans*-Pinocarveol** | 13.48 | 19.16 | **<0.001** | 0.41±0.11(b) | 0.85±0.22(b) | 1.29±0.34(ab) | 2.06±0.31(a) |
| **Camphor** | 13.63 | 12 | **0.003** | 10.28±1.51(b) | 14.12±1.78(ab) | 16.41±1.98(ab) | 30.9±8.3(a) |
| Camphene hydrate | 13.73 | 2.57 | 0.093 | 0.18±0.03 | 0.25±0.05 | 0.31±0.05 | 0.35±0.03 |
| Pinocamphone | 14.43 | 0.13 | 0.727 | 4.81±0.62 | 4.11±0.73 | 3±0.56 | 3.03±0.61 |
| **Pinocarvone** | 14.10 | 14.48 | **0.003** | 0.29±0.04(a) | 0.22±0.08(ab) | 0.05±0.01(ab) | 0.03±0.01(b) |
| *endo*-Borneol | 14.18 | 1.29 | 0.275 | 1.77±0.41 | 3.34±0.51 | 4.18±0.61 | 9.9±1.13 |
| 3-Thujene-2-one | 14.34 | - | - | ND | ND | ND | ND |
| Isopinocamphone | 14.40 | 1.84 | 0.193 | 3.08±0.95 | 4.14±1.56 | 4.57±1.32 | 6.33±2.07 |
| Terpinen-4-ol | 14.46 | 0 | 0.988 | 1.18±0.13 | 1.51±0.34 | 1.37±0.32 | 1.5±0.38 |
| ***p*-Cymene-8-ol** | 14.65 | 178.59 | **<0.001** | 0.36±0.05(d) | 0.8±0.06(c) | 1.33±0.12(b) | 1.97±0.09(a) |
| **α-Terpineol** | 14.79 | 9.08 | **0.008** | 2.97±0.43(b) | 3.77±0.11(ab) | 3.97±0.51(ab) | 5.09±0.6(a) |
| **Myrtenol** | 14.94 | 53.54 | **<0.001** | 0.32±0.08(c) | 2±0.49(b) | 4.44±0.99(ab) | 9.5±2.02(a) |
| Verbenone | 15.28 | 2.85 | 0.126 | 0.19±0.02 | 0.22±0.01 | 0.45±0.13 | 0.35±0.07 |
| 2-Hydroxycineole | 15.58 | 1.04 | 0.332 | ND | 0.09±0 | 0.1±0.03 | 0.15±0.05 |
| **Thymol methyl ether** | 15.85 | 18.59 | **<0.001** | 2.23±0.36(a) | 1.84±0.27(ab) | 1.03±0.25(bc) | 0.83±0.14(c) |
| Myrtanol isomer1 | 16.73 | - | - | ND | ND | TR | TR |
| Myrtanol isomer2 | 16.30 | - | - | TR | TR | TR | 0.1±0.03 |
| *p*-Menth-2-en-7-ol | 16.42 | - | - | ND | ND | ND | ND |
| Myrtanol isomer3 | 16.48 | 2 | 0.2 | ND | 0.05±0(a) | 0.14±0.03(ab) | 0.28±0.11(b) |
| **Myrtenyl acetate isomer1** | 17.43 | 5.73 | **0.048** | 1.15±0.49 | 0.34±0.01 | 0.1±0.02 | ND |
| Myrtenyl acetate isomer2 | 18.13 | - | - | ND | ND | ND | ND |
| ***Sesquiterpenes*** |  | | | | | | |
| **α-Longipinene** | 18.63 | 49.99 | **<0.001** | 0.46±0.11(a) | 0.22±0.06(a) | 0.08±0.03(b) | 0.04±0(b) |
| **Longicyclene** | 19.10 | 40.08 | **<0.001** | 0.52±0.12(a) | 0.28±0.07(a) | 0.09±0.04(b) | 0.05±0(b) |
| Longifolene | 19.88 | 17.97 | 0.001 | 3.23±0.92(a) | 2.23±0.42(ab) | 1.1±0.36(bc) | 0.8±0.19(c) |
| **(*E*)-β-Caryophyllene** | 20.17 | 30.57 | **<0.001** | 10.3±2.52(a) | 6.87±1.39(ab) | 2.95±0.85(bc) | 1.94±0.46(c) |
| (*E*)-β-Caryophyllene (fungus) | 20.56 | - | - | ND | ND | ND | ND |
| (*E*)-β-Farnesene | 20.84 | 3.44 | 0.106 | 0.58±0.16 | 0.48±0.05 | 0.16±0.05 | ND |
| **Humulene** | 20.90 | 18.26 | **0.001** | 3.94±0.86(a) | 2.23±0.41(ab) | 1.38±0.48(b) | 1.02±0.26(b) |
| Caryophyllene oxide | 23.56 | - | - | ND | ND | ND | ND |

^#^- Estimated retention time from GC-MS

***^$^-***Significant differences between time points are denoted by small letters (ANOVA, followed by Tukey’s test, *P<0.05*

| **Compounds** | **Purity^$^** | | **CAS number** | | **Composition (V/V %)** |
| --- | --- | --- | --- | --- | --- |
| Tricyclene | >99 | 508-32-7 | | 0.33 | |
| α-Thujene | 80 | 2867-05-2 | | 0.27 | |
| (-)-α-Pinene | >99 | 80-56-8 | | 24.35 | |
| (+)-α-Pinene | >99 | 80-56-8 | | 28.86 | |
| (-)-Camphene | >99 | 79-92-5 | | 0.98 | |
| (+)-Camphene | >99 | 79-92-5 | | 0.33 | |
| (-)-Sabinene | 76 | 3387-41-5 | | 0.38 | |
| (+)-Sabinene | 76 | 3387-41-5 | | 1.85 | |
| (-)-β-Pinene | >99 | 127-91-3 | | 34.18 | |
| Myrcene | 93 | 123-35-3 | | 2.93 | |
| α-Phellandrene | >99 | 99-83-2 | | 0.16 | |
| *delta*-3-Carene | >99 | 13466-78-9 | | 1.47 | |
| α-Terpinene | 92 | 99-86-5 | | 0.16 | |
| *p*-Cymene | >99 | 99-87-6 | | 0.33 | |
| (-)-Limonene | >99 | 5989-27 | | 0.82 | |
| (+)-Limonene | >99 | 5989-27 | | 0.82 | |
| 1,8-Cineole | >99 | 470-82-6 | | 0.16 | |
| γ-Terpinene | 97 | 99-86-5 | | 0.33 | |
| Terpinolene | >99 | 586-62-9 | | 0.65 | |
| (-)-Bornyl acetate | >99 | 76-49-3 | | 0.65 | |

***Table S8*:** Composition of synthetic monoterpene mixture used in bioassays. ^$^ The purity of each compound was calculated from GC-MS analysis.

|  |  | Colony forming units (CFUs)/mL | | | | | |
| --- | --- | --- | --- | --- | --- | --- | --- |
| Beetle treatment | Sample type | Potato dextrose agar | | | Luria agar | | |
|  |  | *Bacteria* | *Yeast* | *Ophiostomatoid fungi* | *Bacteria* | *Yeast* | *Ophiosotmatoid fungi* |
| Unaltered | Wash | NP | 6083 ± 2655 | NP | NP | 6166 ± 2565 | NP |
|  | Lysate | 1167 ± 385 | 500 ± 236 | 250 ± 156 | 2167 ± 481 | 917 ± 478 | 83 ± 76 |
| Fungus-free (FF) | Wash | NP | NP | NP | NP | NP | NP |
|  | Lysate | >10^5 | 2100 ± 1565 | NP | >10^5 | NP | NP |
| *G. penicillata*-reinoculated in FF | Wash | 900 ± 415 | NP | 340 ± 154 | 900 ± 638 | NP | 300 ± 163 |
|  | Lysate | 62000 ± 33367 | NP | 100 ± 75 | 39000 ± 23749 | NP | 100 ± 81 |

***Table S9*:** Average colony forming units (CFUs/mL) from untreated, fungus-free, and fungus-free, *G. penicillata*-reinoculated *I. typographus* bark beetles (*n* = 5 or 6 beetles). Wash, supernatant from beetles immersed in 0.05 % Triton X in 500 µL PBS buffer pH 7.4; lysate, crushed beetles in 500 µL PBS buffer pH 7.4; NP, not present

| Treatments | Colony forming units (CFUs)/mL | | | |
| --- | --- | --- | --- | --- |
|  | *Bacteria* | *Yeasts* | *Ophiostomatoid fungi* | *Molds/other fungi* |
| Fungus-free | >10^6 | NP | NP | NP |
| Fungus-free | >10^6 | 20 | NP | 2000 (red mold), 60 (dark green mold) |
| Fungus-free | >10^6 | 30 | NP | 40 (dark green mold) |
| Fungus-free | 70 | NP | NP | NP |
| Fungus-free | NP | 210 | NP | 140 (dark green mold), 3000 (red mold) |
| Unaltered | NP | 27000 | 70 | NP |
| Unaltered | 9000 | 55000 | 10 | NP |
| Unaltered | 21000 | 3000 | 40 | NP |
| Unaltered | NP | NP | 10 | 1200 (red mold) |
| Unaltered | NP | 21000 | 20 | NP |
| Unaltered | NP | 13000 | 10 | NP |
| Unaltered | >10^6 | 272000 | NP | NP |
| *G. penicillata*-reinoculated FF^$^ | NP | NP | 100 | NP |

***Table S10*:** Colony forming units (CFUs/mL) obtained from bark beetle gallery samples infested by fungus-free beetles, and fungus-free beetles reinoculated with *G. penicillata*, and untreated control beetles. Approximately 300 mg of bark samples were dissolved in 1 mL PBS buffer solution and dilutions were plated on PDA. NP, not present;

^$^ Only one gallery sample was tested due to low sample availability.
