## Supplemental methods for "Bark beetles locate fungal symbionts by detecting volatile fungal metabolites of host tree resin monoterpenes"

**Preparation of bark beetle diet for eliminating fungal symbionts**

This semi-artificial diet was adapted from [1–4] and contains 30 % freeze-dried finely powdered spruce phloem, 6 % inactivated yeast, 1 % sodium benzoate, and 63 % water. Approximately 2 g of mixed diet was added to 19*65mm shell vials (Kimble, VWR) using a 10 ml plastic syringe. The mouth of the syringe was cut so that the diameter at the tip of the syringe was same as its inner diameter. The plunger inside the syringe was pulled to create space around 3 ml volume and with the help of spatula, the diet mixture was added into the syringe by pressing gently using the spatula. The diet was then packed into shell vials by inserting the syringe containing the diet to reach the bottom of the vial and then the diet was dispensed carefully by pressing the plunger to avoid contaminating the walls of the vial with diet particles. To achieve the texture preferred by beetles, the diet mixture added in the vials was pressed gently until the diet was tightly packed. Using 2 mm sterile plastic rod, a small hole of approximately 8 mm height was made in each vial. A single pupa of *I. typographus* was carefully added into each hole with its head facing towards the diet using a wet paint brush. Pupae were obtained as follows: fresh spruce logs were infested with parent beetles and rearing conditions were set at constant temperature 23°C, relative humidity 65% and photoperiod 18L:6D. After 22-25 days, bark of the breeding logs was peeled to reveal beetle galleries, and white pupae were collected using a wet paint brush and kept in 2 % water agar until used.

After adding pupae, diet vials were kept in the rearing chamber under the same conditions as mentioned above. Approximately after 10 days, fungus-free, mature beetles were collected from the diet vials. Adult beetles when they attain maturation, they emerge and stay on top of the diet. To reinfect ophiostomatoid fungus in beetles, *G. penicillata* was inoculated in the same diet used for beetles except sodium benzoate was replaced by 0.05 % cycloheximide to which *G. penicillata* is resistant. Fungus-free adult beetles were then placed in *G. penicillata*-inoculated diet vials and allowed to tunnel for 1 day. 1) Sterility of the diet, 2) sterility of beetles reared on the semi-artificial diet for fungal elimination, and 3) reinfection of *G. penicillata* in fungus-free adults were verified by plating beetle frass or crushed beetles on PDA and also by imaging beetle parts using scanning electron microscopy (SEM). This protocol provided fungus-free mature beetles from pupae and fungus-free beetles reinfected with *G. penicillata* (S9 Table, Fig. 3A).

Fungus-free beetles, *G. penicillata*-reinoculated beetles, and untreated control beetles with their usual complement of fungi were infested in fresh spruce logs. Untreated beetles were obtained from laboratory culture (generation 9) maintained at the rearing condition mentioned above; the culture originated from Asa, Sweden. Two logs were used for each treatment and 3 pairs of treated beetles (fungus-free and *G. penicillata*-reinoculated beetles), and 5 pairs of untreated beetles were used to infest their corresponding logs. After 10 days, bark of the logs was peeled, and the beetle gallery samples were flash frozen in liquid N_2_ for terpene analysis and stored at 4°C for microbial test. Beetle-free and discolored bark samples were taken from each log at the end of experiment for control chemical analysis.

**Analysis of *G. penicillata* and *Trichoderma sp.* headspace volatiles**

Headspace volatiles from *G. penicillata* and *Trichoderma sp.* grown on PDA amended with terpene mix were collected using SPME fiber (coating-50/30 µm DVB/CAR/PDMS; core/assembly type-StableFlex^TM^/ stainless steel (1cm), Supelco). Briefly, 1 ml of PDA containing 0.1 mg/ml of mix of terpenes was poured in 15 ml glass test tubes and allowed to solidify before *G. penicillata* and Trichoderma sp. were inoculated. Control tubes contained fungus-free PDA plus terpenes. After 5 d post inoculation, headspace volatiles were collected for 15 mins from each test tube using SPME fiber and were manually injected immediately in the GC inlet. The fibers were conditioned prior to headspace collection and cleaned in between each collection following the manufacturer’s protocol. GC-MS analysis was carried out in Agilent 7890A GC system coupled with Agilent 5975C mass detector. Injection program was configured in splitless mode at 250°C. Non-polar HP-5MS GC column (30m *250µm* 0.25 µm) was used to separate compounds and helium was used as a carrier gas. The oven temperature started at 45°C and held for 1 min and then increased to 250°C at the rate of 10°C min^-1^ and finally held for 5 min. Compounds were identified by comparing their mass spectra with NIST reference library and later by matching their retention time and mass spectra to those of authentic standards.

**Beetle pheromone analysis**

For identification of terpene metabolites produced by adult bark beetles, beetle fumigation experiment was carried out using synthetic terpene mixture in 20 ml scintillation vials. Briefly, two males were placed in clean 20 ml glass vials and 2 µl of terpene mixture was added to a clean 23 mm circular filter paper (Whatmann no. 3) that was secured tightly under the screw cap. Beetles were not in the direct contact with terpenes, but they were exposed to terpene vapors. Control treatments were dead beetles (freeze-killed) exposed to terpene vapors and live beetles without terpene vapors in glass vials. Each treatment was replicated 8 times. Terpene metabolites from beetles were extracted as follows: four beetles from each treatment were placed in 4 ml glass vials containing 1 ml of heptane and crushed with a clean plastic pestle. Then the slurry was centrifuged at 4000 rpm for 5 min and supernatant were transferred to a clean 1.5 ml glass vials. Since the beetle extract contained high concentration of fatty acids, the extract was fractionated to separate fatty acids from polar compounds. Glass capillary was packed with 400 mg of Florosil^®^ (Sigma-Aldrich) and conditioned with 2 ml heptane. Beetle extract was concentrated to 100 µL under N_2_ air and added to the column. Then the vial that contained beetle extract was rinsed twice with 100 µL to remove the left-over extract and added to the same column. The column was first washed with 2 mL heptane to remove all the hydrocarbons. Second wash was performed with 2 ml heptane + 20% acetone to collect all oxygenated derivatives that include terpenoids. Final wash was done with 2 mL heptane + 50 % acetone to remove left-over polar compounds and conditioned again with 2 ml heptane before the next beetle extract was loaded for fractionation. Each fraction was analyzed in GC-MS to identify terpenoids produced by beetles and the concentration in each fraction was quantified using external calibration curves. The GC program was same as described before and injection volume was set at 2 µL.

1.	Colineau B, Lieutier F. Production of Ophiostoma-free adults *of Ips sexdentatus* boern. (Coleoptera: Scolytidae) and comparison with naturally contaminated adults. Can Entomol. 1994;126: 103–110. doi:10.4039/Ent126103-1

2.	Schmidt FH. Two Artificial (Oligidic) Media for the Douglas-fir Beetle, *Dendroctonus pseudotsugae* Hopkins (Coleoptera: Scolytidae). CanEntomol. 1966;98: 1050–1055. doi:10.4039/Ent981050-10

3.	Whitney HS, Spanier OJ. An improved method for rearing axenic mountain pine beetles, *Dendroctonus* *ponderosae* (Coleoptera: Scolytidae). Can Entomol. 1982;114: 1095–1100. doi:10.4039/Ent1141095-11

4.	Bedard WD. A ground phloem medium for rearing immature bark beetles (Scolytidae). Ann Entomol Soc Am. 1966;59: 931–938. doi:10.1093/aesa/59.5.931
